# Reproducing extracellular matrix adverse remodelling of non-ST myocardial infarction in a large animal model

**DOI:** 10.1101/2022.05.19.492645

**Authors:** Paolo Contessotto, Renza Spelat, Vaidas Vysockas, Aušra Krivickienė, Chunsheng Jin, Sandrine Chantepie, Clizia Chinello, Audrys G. Pauza, Mindaugas Rackauskas, Vilma Zigmantaitė, Fulvio Magni, Dulce Papy-Garcia, Niclas G. Karlsson, Eglė Ereminienė, Abhay Pandit, Mark Da Costa

## Abstract

The rising incidence of non-ST-segment elevation myocardial infarction (NSTEMI) and associated long-term high mortality constitutes an urgent clinical issue. Unfortunately, the study of possible interventions to treat this pathology lacks a reproducible pre-clinical model. Indeed, currently adopted small and large animal models of MI mimic only full-thickness, ST-segment-elevation (STEMI) infarcts, and hence cater only for investigation into therapeutics and interventions directed at this subset of MI. Thus, we developed an ovine model of NSTEMI by ligating the myocardial muscle at precise intervals parallel to the left anterior descending coronary artery. After validating the presented model both by histology and functional analysis with clinical data, further omics analyses highlighted the distinctive features of post-NSTEMI tissue remodelling. Here, by looking at the transcriptome and proteome-derived pathways emerging at acute (7 days) and late (28 days) post-surgery timepoints, we discovered specific alterations in cardiac post-ischaemic extracellular matrix (ECM). Together with the rise of well-known markers of inflammation and fibrosis, NSTEMI ischaemic regions showed distinctive patterns in the expression of complex N-glycans and glycosaminoglycans in cellular membranes and ECM. Identifying such changes in molecular moieties accessible to infusible and intra-myocardial injectable drugs sheds light on the development of targeted pharmacological solutions to contrast adverse fibrotic remodelling.

---

Myocardial infarction (MI) belongs to the family of coronary artery diseases and is the leading cause of cardiovascular-related worldwide mortality^1^. In addition, COVID-19 infection was recently shown to be an additional major risk factor in non-hospitalised cases^2^. Patients who survive an MI often suffer lethal heart failure later on. Indeed, heart failure appears to be an influential adverse prognostic factor after an MI^3^. A marked rise in non-ST-elevation myocardial infarctions (NSTEMIs) in hospitalised cases has emerged over the last two decades^4,5^. Moreover, despite the smaller cardiac ventricular areas affected by the ischaemic event, registry data show that NSTEMIs are associated with long-term mortality that is higher than that after ST-elevation myocardial infarctions (STEMIs)^6-9^.

In current preclinical studies MI is mainly reproduced as STEMI both in rodents (rats and murine) and large animals (porcine and ovine) by the ligation of the coronary arteries arising from the left coronary artery, specifically with an established preference for the ligation of the left anterior descending coronary artery (LAD)^10,11^. Nonetheless, ligation of the LAD results in an extended infarct in the left ventricle, associated with high experimental mortality and a poor reflection of most hospitalised clinical cases. Indeed, most MIs currently reported in the clinics are either partial or non-transmural infarcts, and often involve multiple small regions of the left ventricle^5,12^. Only a few studies have paved the way by optimising the extension of the infarct up to approximately 25% of the infarct’s left ventricular mass and bringing the overall experimental mortality down to around 17%^13,14^. Therefore, there is a clear need in the field to adopt validated, clinically similar models to evaluate NSTEMI pathophysiology and possible interventions fully.

In all the previously adopted preclinical models of MI, collagen deposition in the myocardial left ventricular wall concludes the process which starts with the ischaemic insult. Fibrosis compensates, albeit poorly, for the extensive loss of cardiomyocytes^15^. Indeed, the entire post-infarction process begins with a sterile immunological response involving different populations of macrophages and inflammatory cells that have only recently been characterised by single-cell RNA sequencing^16,17^. Significantly, post-ischaemic myocardial remodelling disrupts the initial balance of glycosaminoglycans (GAGs), proteoglycans and glycans which are present in the extracellular matrix (ECM) and cell membrane of cardiac cell populations (cardiomyocytes, fibroblasts, endothelial cells)^18,19^. Consequently, adverse remodelling affects the structural and mechanical stability of the ECM environment. This also leads to imbalances in molecular pathways that include the recruitment of growth factors such as vascular endothelial growth factor (VEGF), platelet-derived growth factor (PDGF) and fibroblast growth factor (FGF). Most studies on MI are performed by reproducing full-thickness STEMIs in mice and rodents, and this is an obvious limitation in their translation to humans. Large animals, specifically sheep, have been extensively used to evaluate the recovery of heart functionality following STEMI because of the similarity in the organ volume with that in humans^20,21^.

In the proposed ovine model of NSTEMI we have analysed the molecular and histological features both at an acute (7 days) timepoint during repair and healing, and a late (28 days) timepoint post-MI. Indeed, given the key role of the modulation of post-ischaemic cardiac ECM in developing translational therapies and the lack of an established animal model that mimics partial thickness myocardial infarction, we felt that a clinically relevant translational model of NSTEMI was urgently needed in the cardiovascular field^4,22,23^. Once we established a surgical procedure to achieve clinically similar NSTEMIs in sheep, we investigated the ischaemic, border zone and remote regions at early (7 days) and late (28 days) timepoints post-MI by histology, RNA-sequencing, proteomics and glycomics. We hope that the proposed NSTEMI model would inspire specific translational options to target this world-wide increasing pathology.

## Results

### Stable functional impairment in a clinically relevant ovine model of NSTEMI

Current preclinical models of full-thickness infarcts (STEMIs) are based on the ligation of the LAD or variations of this procedure in the LAD territory^10,24^. A significant limitation of the LAD ligation model in its clinical resemblance is that proximal occlusion of the coronary is often fatal. Indeed, the incidence of NSTEMIs currently exceeds the number of actual clinical infarcts that LAD ligation-based models aim to reproduce^22,25,26^. In this study, multiple ligations were performed lateral and parallel to the LAD from the first diagonal up to 3-4 cm from the apex to induce multiple non-transmural infarcts in the left ventricle (Fig. 1a,b and Extended Data Fig. 1). The proposed model of NSTEMI was performed in a cohort of 21 sheep. Four sheep died during the surgical procedure, resulting in an overall mortality of 19.04%. Vagus nerve stimulation during intubation caused two deaths, one was due to left ventricle rupture and one of uncertain cause at the premedication stage. Thus, only one sheep died from the actual MI, specifically from a complication of the induced MI. Six sheep were sacrificed on day 7 (d7) post-MI and the remaining eleven on day 28 (d28) post-NSTEMI as the final endpoint to evaluate both functional and histological alterations.

**Figure 1.**
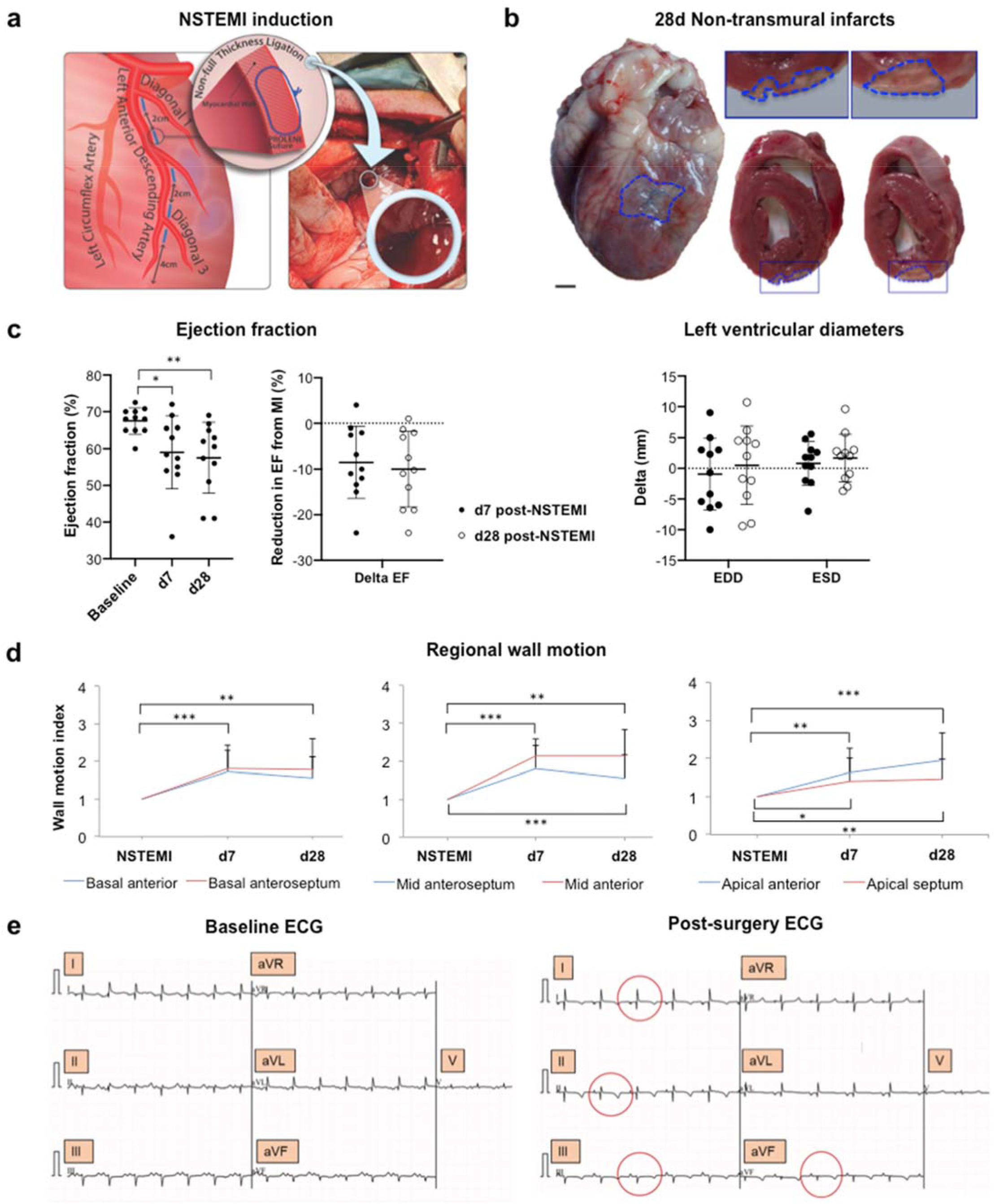
Clinically relevant ovine model of NSTEMI. **a**, Schematics of the multiple ligation procedure to induce NSTEMI infarcts. **b**, Representative photographs of 8-month-old explanted and axially-cut sheep hearts 28 days post-ligation. Blue Prolene sutures were used to track NSTEMI infarcts (framed in blue). *n*=11 animals. 1-mm ruler spacing. Insets shown at higher magnification above. Scale bar, 1 cm. **c**, Left, ejection fraction (EF) absolute values before ligation (baseline), 7 (d7) and 28 (d28) days post-ligation and relative decrease in EF on d7 and d28 post-surgery (left). Right, measurement of left ventricular end diastolic (EDD) and systolic (ESD) on d7 and 28 post-ligation. *n*=11 animals. **d**, Regional wall motion analysis in the main six cardiac segments affected by the induction of NSTEMI by ligation. Wall motion index is shown as mean ± s.d at d7 and d28 post-NSTEMI induction. *n*=11 animals. **e**, Representative electrocardiogram (ECG) before NSTEMI-induction (left) and post-ligation (right). Changes in T wave inversion, in leads I, II, III and aVF are circled in red. *n*=4 animals. Kruskal-Wallis test in (**c**), multiple unpaired t-test with Benjamini’s method in (**d**). \**P*<0.05, \*\**P*<0.01, \*\*\**P*<0.001.

Full-thickness infarcts induce marked drops in ejection fraction (EF)^27,28^. In contrast, the current NSTEMI model causes focal infarcts, associated with a limited yet significant reduction in EF (8.52±7.88%, P=0.03) on d7 post-surgery (Fig. 1c). Three weeks later (d28), EF decreased by 10±8.31% (P=0.009) relative to the pre-MI levels (Fig. 1c). Post-ischaemic remodelling involves different degrees of dilatation, hypertrophy and finally collagen scarring. This process occurs over weeks and months and it is influenced by multiple factors including the size and site of the infarct, whether the infarct was transmural (STEMI) or not (NSTEMI), the amount of stunning of the peri-infarct myocardium, the patency of the related coronary artery and local trophic factors^29,30^. Since NSTEMI is not the result of complete occlusion of a coronary artery, it usually affects a small area or those that are diffuse or patchy areas of the ventricular muscle rather than the entire thickness of the local ventricular wall. Indeed, the presented model did not significantly vary in left ventricular diastolic and systolic diameters (LVEDD and LVESD) (Fig. 1c). Therefore, to further evaluate the functional impairment seen in the proposed NSTEMI model, we analysed the loss of cardiac contractility in the specific left ventricular segments affected by NSTEMI through regional wall motion index (WMI) analysis (Fig. 1d). On d7 post-NSTEMI wall motion impairments were significant in basal anterior/anteroseptum, (WMI=1.73±0.56 and 1.82±0.6, P<0.001), mid anteroseptum/anterior (WMI=1.82±0.6 and 2.14±0.45, P<0.001), apical anterior (WMI=1.64±0.64, P<0.001), and apical septum (WMI=1.41±0.58, P<0.05) segments. This widespread wall-motion deficit persisted in all the affected segments on d28 post-NSTEMI (Fig. 1d).

To validate the current NSTEMI induction with clinical data, electrocardiograms (ECGs) post-ligation highlighted comparable changes in T wave inversion in leads I, II, III and aVF (Fig. 1e and Extended Data Fig. 2). In line with presented data on LVEDD and LVESD, the reduction in fractional shortening (FS) was not significant on d7 and d28 post-surgery (Extended Data Fig. 3a). In addition, we measured a significant rise in troponin I serum levels on day 1 and 2 post-NSTEMI induction (Extended Data Fig. 3b). This data further supports the required criteria to define the current model as representative of an NSTEMI^31^.

### Ischaemic damage and consequent adverse remodelling following NSTEMI

The non-transmural nature of the induced NSTEMI infarcts became apparent with explantation on d28 post-MI (Fig. 1b). During the surgical procedure, the localisation of the infarct was defined by its proximity to the blue suture used to perform the multiple ligations (Extended Data Fig. 1b-f). Indeed, once the hearts were sliced at a thickness of 1 cm, the NSTEMI regions were detectable by the discolouration of the left ventricular wall (Fig. 1b). Therefore, sampling was carried out from the core ischaemic progressively to the border and remote regions (Extended Data Fig. 3c). Specimens were isotropically uniformly oriented to avoid any bias when evaluating cardiomyocytes and vasculature structures^32^. It is well known that the necrotic phase responsible for the loss of cardiomyocytes in the left ventricle and the consequent highly pro-inflammatory microenvironment represents the first immediate steps post-ischaemia^33^. Moreover, a valid model of MI also needs to reproduce the long-term effects on the border zone regions indirectly affected by the complete tissue disruption in the ischaemic core^34,35^. This study detected a progressive vacuolisation inside the cardiomyocytes from d7 to d28 after NSTEMI induction (Fig. 2a). Disruption of the intercalated disks and lipid droplet accumulation (Fig. 2a and Extended Data Fig. 3d) were seen exclusively in the acute phase (d7). In contrast, fibrotic deposition fully developed at the endpoint (d28) of the study (Fig. 2a,b and Extended Data Fig. 3d). Specifically, intercalated disk disruption highlighted the structural disassembly of the cardiomyocyte myofibril apparatus needed for its cellular contraction and thus the synchronous beating of the whole organ. In addition, the accumulation of dense bodies in the mitochondria of these cardiomyocytes further supported the evidence of ischaemic injury (Fig. 2c). Together, these data show that the current model of NSTEMI reproduced the sequential inflammatory and fibrotic remodelling of the infarcted region, resulting in a non-full-thickness scar formation which is characteristic of the clinical cases of NSTEMI. Histology showed the extended loss of cardiomyocytes from the ischaemic core to the border zone on d7 post-NSTEMI induction (Fig. 2b). From the end of the first week after MI and through the following weeks the myofibroblasts secrete collagen^36^, and this event was also observed in the current model (Fig. 2b,d). Moreover, an irregular vascularisation in the fibrotic areas was detected by immunostaining for α-SMA^+^ arterioles. (Fig. 2d).

**Figure 2.**
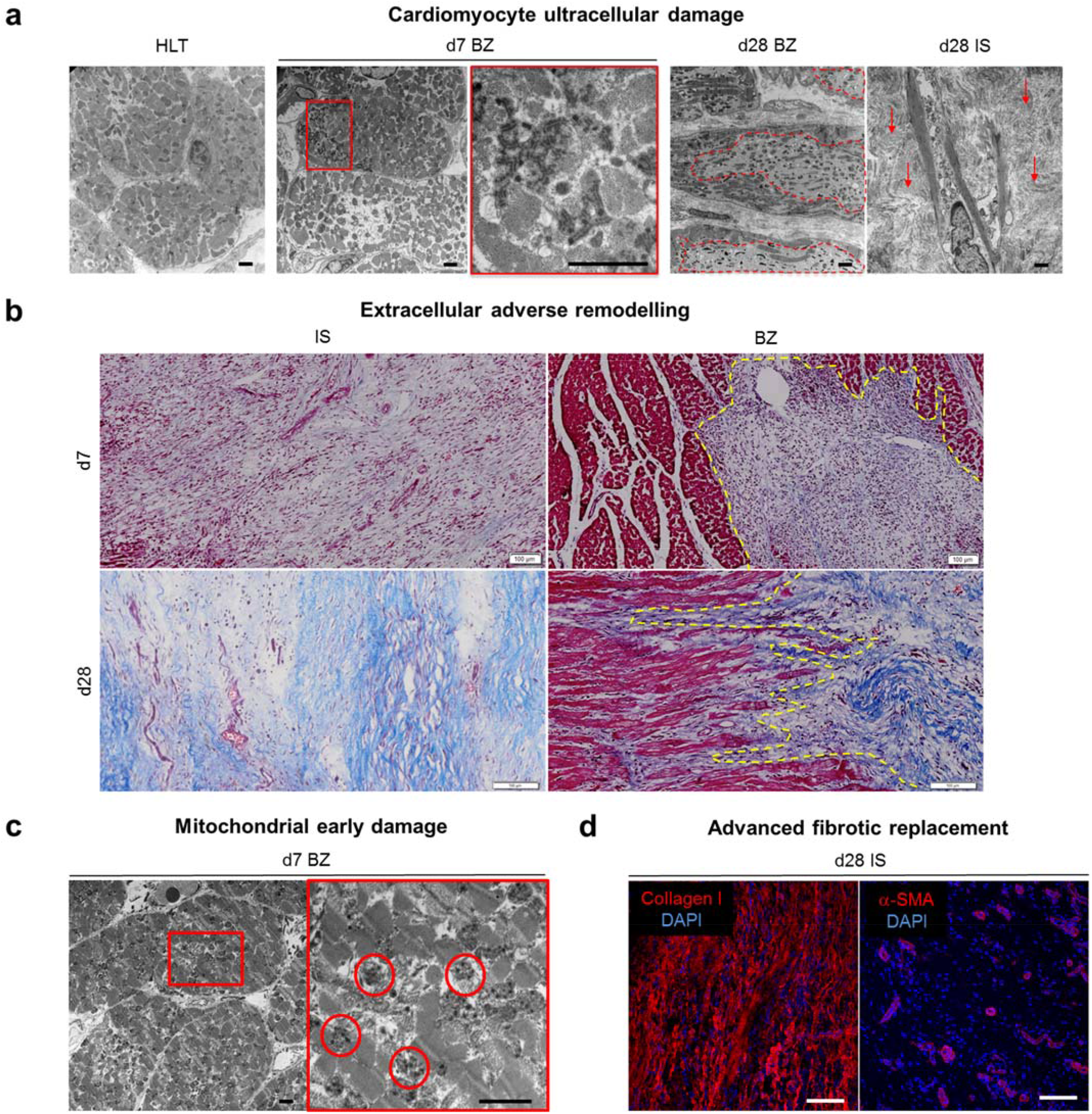
Ischaemic cellular damage and extracellular adverse remodelling following NSTEMI. **a**, Representative TEM micrographs showing ultracellular damage in healthy (HLT) cardiomyocytes (far left) starting from intercalated disks disruption on d7 post-NSTEMI (center), to extended vacuolisation (dashed red line) on d28 post-NSTEMI (right), surrounded by collagen deposition (arrows) by myofibrobalsts. *n*=5 HLT and d7, *n*=7 d28 animals. Scale bars, 2 µm. **b**, Representative Masson’s Trichrome staining of ischemic core (IS) and border zone (BZ) regions of NSTEMI infarcted tissues on d7 and d28. *n*=5 animals per group. Scale bars, 100 µm. **c**, Representative TEM micrographs of mitochondria in cardiomyocytes located in the BZ of the infarct. Inset shows accumulation of dense bodies (circled in red) on d7 post-NSTEMI. *n*=5 animals. Scale bars, 2 μm. **d**, Immunofluorescence microscopy of collagen fibrotic replacement (left) and sparse α-SMA^+^ arterioles (right) in IS on d28. *n*=5 animals. Scale bars, 20 μm.

### Transcriptome and proteome of NSTEMI infarcts

To investigate the molecular profiling of the current model of NSTEMI, we analysed the transcriptome and proteome of the ischaemic core, border, and remote region. Bulk RNA-sequencing analysis (adjusted p value of 0.05, mean quality score 38.41) highlighted pools of differentially expressed genes (DEG) at each timepoint post-ligation, depending on the distance from the ischaemic core (Fig. 3a). A clear regional- and temporal-dependent transcriptome alteration was seen on the ischaemic core, border and remote regions (Fig. 3a and Extended Data Fig. 4). Specifically, 1079 transcripts were upregulated (log2Fold-change>1.5) and 805 downregulated (log2Fold-change<1.5) in the ischaemic core on d7 compared to d28 post-NSTEMI (Fig. 3a). Moreover, in the border zone 1855 transcripts were upregulated and 667 downregulated, and finally in the remote region 284 were increased and 233 decreased on d7 compared to d28 post-surgery (Fig. 3a and Extended Data Fig. 4).

**Figure 3.**
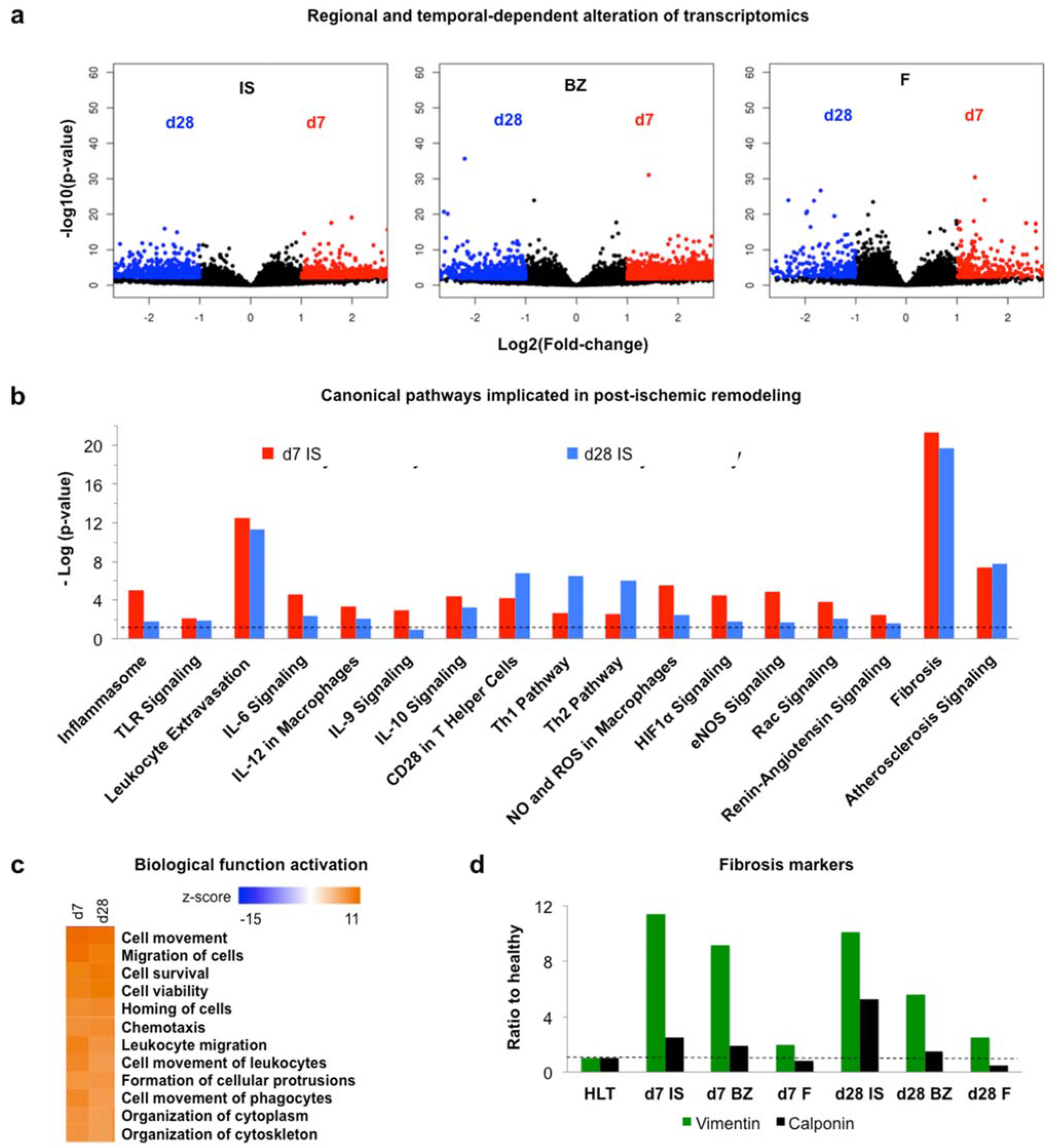
Post-ischaemic pathways alteration following NSTEMI. **a**, Volcano plots showing the total genes significantly upregulated (log2(fold change) > 1.5) between the core ischaemic (IS), border (BZ) and remote (F) regions sampled at d7 and d28 post-NSTEMI. *n*=4 animals per group. **b**, Significant canonical pathways resulting from differentially expressed genes (DEG) data from RNA sequencing (analysed by IPA^®^). Cut-offs of log2(fold change) > 1.5 and log2(fold change) < -1.5 and adjusted-*P*<0.05 were set. Dashed line shows a threshold of -Log(p-value) of 1.3, corresponding to *P*=0.05. All DEG data were normalised to healthy baseline left ventricular samples. *n*=4 animals per group. **c**, Main activated biological functions listed by highest z-score from IPA^®^ analysis on RNA-seq data from IS samples on d7 and d28 post-NSTEMI. *n*=4 animals per group. **d**, Expression levels of myofibroblast-related markers vimentin and calponin 1 as detected by nLC-ESI-MS/MS analysis on IS samples. Each analysed sample was a pool of samples coming from three animals.

DEG data derived from infarcted tissues on d7 and 28 vs healthy samples were analysed by Ingenuity Pathway Analysis (IPA^®^) to identify which canonical pathways are significantly altered following NSTEMI. Many typical pathways associated with myocardial ischaemic pathophysiology were detected, such as fibrosis and atherosclerotic, renin-angiotensin signalling both on d7 and 28 post-surgery (Fig. 3b). In addition, several pathways linked to the inflammatory response emerged in the d7 post-NSTEMI group, including hypoxia-inducible factor (HIF1α), endothelial NOS (eNOS), and interleukin-6 (IL-6), -9, -10, -12, leukocyte extravasation and inflammasome signalling (Fig. 3b). On d7 and d28 post-NSTEMI biological functions linked to inflammatory cell recruitment ranked as the top activated ones (z-score above 10) (Fig. 3c). Indeed, pathway analysis on DEG associated with higher HIF1α expression predicted the activation of inflammatory markers such as IL-1β, IL-6, and tumour necrosis factor (TNF) on d7 (Extended Data Fig. 5a). HIF1α was also associated with the predicted activation of fibrotic molecular profiling such as transforming growth factor-β (TGF-β), matrix metalloproteinase-2 and -9 (MMP-2, -9) and lysyl oxidase (LOX) on d28 in the ischaemic core (Extended Data Fig. 5b).

Proteomic analysis was run to validate further the molecular changes seen by RNA-seq and pathway analysis showing a post-ischaemic remodelling response. In line with the RNA-seq outcome, pathway analysis on proteomic data showed significant activation of IL1-β, TNF, IL-6, TGF-β, IFN-γ in the ischaemic core 28 days post-NSTEMI (Extended Data Table 1). In addition, LC-ESI-MS/MS data from whole extracts of ischaemic core, border, and remote regions (false discovery rate below 1%) highlighted well-known markers of fibrotic replacement (Fig. 3d). Interestingly, a progressively increased calponin 1 and vimentin expression was seen from the remote to the ischaemic core regions both on d7 and d28 post-NSTEMI (Fig. 3d). Specifically, when compared with the healthy left ventricular myocardial sample, vimentin ratio increased to 9.13-fold and 11.37-fold on d7 and to 5.58-fold and 10.10-fold on d28 in the border and ischaemic core regions, respectively. Also, the calponin-1 ratio increased to 1.87-fold and 2.5-fold on d7 and to 1.5-fold and 5.28-fold on d28 in the border and ischaemic core regions, respectively (Fig. 3d). Moreover, gene-annotation enrichment analysis using Database for Annotation, Visualization, and Integrated Discovery (DAVID) software on RNA-seq data highlighted glycan alterations in the post-ischaemic remodelling in the current NSTEMI model (Extended Data Fig. 6). Both on d7 post-surgery N-glycan biosynthesis (Extended Data Fig. 6a) and on d28 post-surgery glycosaminoglycans (GAGs) biosynthesis (Extended Data Fig. 6b) emerged among the biological categories (KEGG pathways) with the highest enrichment score. As the relevance of glycoproteins in the pathophysiology of MI has raised increasing interest in recent years^18,37^, we have expanded this finding through advanced glycomics on N-linked glycans extracted from the cellular membrane and ECM proteins in ischaemic, border and remote regions.

### Distinct glycoprofile in the ischaemic tissue following NSTEMI

Considering the findings highlighted by the functional annotation analysis on RNA-seq data, we scrutinised the altered glycan composition in the cellular membrane and ECM proteins of infarcted NSTEMI tissue. Here, we dissected the glycome under the inflammatory (d7) and fibrotic (d28) conditions following NSTEMI by advanced glycomic analysis of the N-glycans expressed in the left ventricular membrane and ECM protein fraction. To achieve this, during the sample processing N-linked glycans were released and analysed by LC-MS. Following annotation procedures, 103 putative N-glycan structures were identified, including 10 high mannose, 15 hybrid and 78 complex-type glycans (Fig. 4). Since the relative abundance of these structures varied across healthy and infarcted tissues, hierarchical clustering analysis was performed for each N-linked glycan subgroup. (Fig. 4a-e). High-mannose N-linked glycans clustering reflected the main distinctions among ischaemic core (IS), border (BZ) and remote regions (F) (Fig. 4a). The subgroup consisting of high mannose N-glycans in the healthy myocardium and F regions showed a global similarity of only 38% to IS on d7 and d28 high mannose N-glycans (based on distribution found by MS) (Fig. 4a). This is in contrast to the post-NSTEMI (d7 and d28), where IS regions showed a global 76% similarity in high mannose N-glycans when grouped. The difference in the level of high mannose glycans showed some interesting features at d7 and d28 post-NSTEMI depending on the distance from the IS region (Fig. 4b). In IS, high mannose structures decreased to 21-24% compared to healthy (HLT) tissue (32%) both on d7 and d28. At the same time, the remote (F) area appeared not to be effected as judged by the similarity in expression compared to healthy. The border zone (BZ) also appeared to be affected by a decrease in high mannose at the early timepoint (d7), but recovered after 28 days, to 38%, a level similar to those of healthy (32%) and less effected remote areas (F) (32-34%) (Fig. 4b).

**Figure 4.**
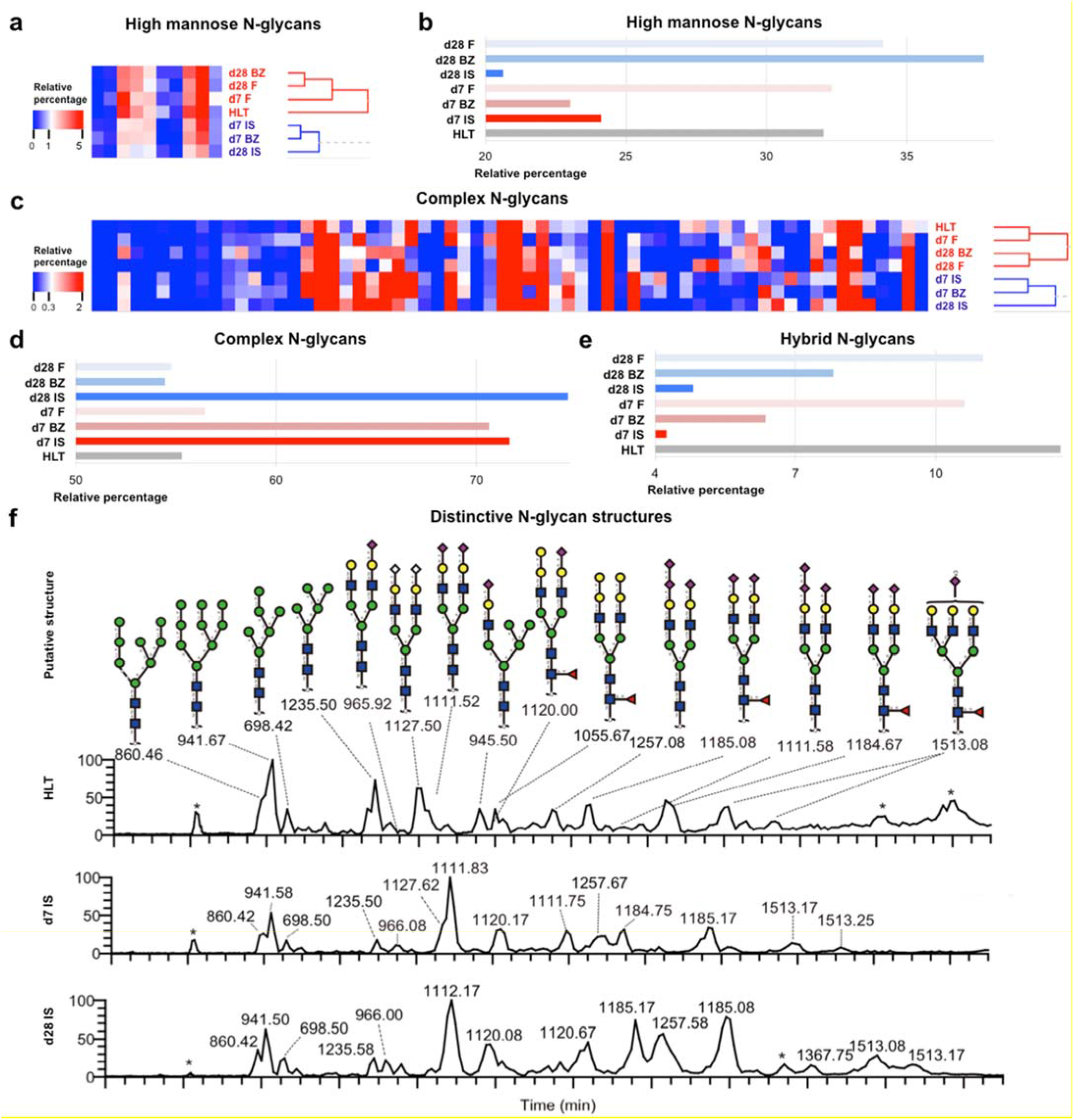
Distinct glycoprofile in the infarcted heart following NSTEMI. **a**, 10 high mannose N-glycans putative structures detected by PG-LC-ESI-MS/MS and analysed by hierarchical clustering. **b**, Relative percentage of high mannose among total N-glycans putative structures across healthy and infarcted myocardial membrane cellular samples at d7 and d28 post-NSTEMI. **c**, 78 complex N-glycans putative structures detected by PG-LC-ESI-MS/MS and analysed by hierarchical clustering. **d**,**e** Relative percentage of complex (**d**) and hybrid (**e**) among total N-glycans putative structures across healthy (HLT) and infarcted myocardial membrane cellular samples at d7 and d28 post-NSTEMI. **f**, Extracted ion chromatography (EIC) showing N-linked glycans mainly expressed in the membrane protein extracts from HLT myocardium and IS at d7 and d28 post-NSTEMI. Regions of infarcted hearts are labelled as follows: IS= core ischaemic, BZ = border zone, F = remote zone from the infarct. Data are representative of two independent experiments. Each analysed sample was a pool of samples coming from three individuals per group and region and analysed by PG-LC-ESI-MS/MS.

The opposite effect, consistent with the trends of the high mannose structure, was seen in complex N-glycans. Indeed, the level of complex N-glycans detected in IS regions on d7 and d28 had a limited similarity (39%) with healthy myocardium (Fig. 4c,d). Ischaemic core regions clustered together both on d7 and d28, and hence showed no signs to recover due to time, with a low (48%) similarity with the less effected remote ones (F) (Fig. 4c). Starting from d7, both in IS and BZ the level of complex glycans was increased (71%) compared to those of healthy and less effected F areas (55%) (Fig. 4d), until d28 in IS (75%). Finally, in IS regions hybrid N-glycan expression decreased to 4-5% from the initial 13% in healthy (Fig. 4e).

Sialylation is a well-known glycosylation type present across cardiac cellular populations such as cardiomyocytes and endothelial cells^18,38^. Given the observed increase in complex N-glycans post-NSTEMI (Fig. 4c,d), we aimed to analyse alterations in sialylated complex N-glycans structures (Fig. 4f). Therefore, we focused on potential differences in the abundance of sialic acid types such as neuraminic acid (NeuAc) and N-glycolylneuraminic acid (NeuGc) compared to healthy cardiac ventricular tissue. NeuAc expression progressively increased from d7 to d28 post-NSTEMI in all infarcted regions, including F, reaching a maximum rise of over 40% in IS on d28 (Extended Data Fig. 7a). The observed NeuGc expression increase on d7 in the infarcted areas (from 34% to 44% more than healthy) was completely lost at the fibrotic timepoint on d28 (Extended Data Fig. 7a). Moreover, we also analysed the sialic acid linkage type observing a progressive increase of α-(2,3)-sialylation in all regions over the remodelling from d7 to d28 (Extended Data Fig. 7b). For the (α-(2,6)-sialic acid linkage, the proportional increase from the healthy state was seen at all timepoints in the IS and BZ areas apart from BZ on d28 (Extended Data Fig. 7b). Finally, a marked difference was seen in the trend of expression of terminal α-gal. The initial marked increase seen on d7 post-NSTEMI in IS and BZ regions (over 30%), was completely lost on d28 in both regions (Extended Data Fig. 7c). We also analysed released O-linked glycans in NSTEMI, identifying the appearance of post-NSTEMI sialylated structures, mainly present during the early phase of remodelling (d7) (Extended Data Fig. 7d).

Altogether, these data indicated a distinct glycoprofile in the NSTEMI cardiac tissue characterised by a higher abundance of complex N-glycans and in particular NeuAc (both (α-(2,3)- and (α-(2,6)-sialic acid linkage) on d7 as well as on d28 post-NSTEMI (Fig. 4f). However, a marked increase in NeuGc and terminal α-galactose (α-gal) was associated only with the early phase of remodelling (d7), but lost through the endpoint (d28) when NeuAc – and in particular α-(2,3)-sialylation – were highly present.

### An irreversibly altered extracellular matrix shows specific changes in the HS sulfation pattern

To further investigate the extensive changes which occur in the ECM following the induction of NSTEMI infarcts, we performed additional analyses on essential components of the myocardial ECM, such as GAGs. Indeed, the restructuring of the ECM is one of the main consequences of post-ischaemic remodelling in the left ventricular wall^33,36^. Specifically, GAGs can regulate inflammation and angiogenesis, influencing the remodelling response^39,40^. The ischaemic core region was initially screened for the presence of sulfated GAGs (sGAGs) by Alcian Blue staining. This confirmed their distribution within the fibrotic regions compared with Masson’s Trichrome staining (Fig. 5a). Then, sGAGs were extracted from samples of the infarct zone harvested 7- and 28-days post-NSTEMI. After total GAGs quantification, the ratio between heparan sulfate (HS) and chondroitin sulfate (CS) was calculated by a subtraction method after enzymatic digestion of the samples with chondroitinase ABC. An increase in CS was seen on d7 post-MI, bringing from 0.98±0.02 in healthy condition to 0.67±0.08, even though this was not significant (p=0.14). Nonetheless, HS/CS ratio significantly (p=0.013) decreased to 0.49±0.13 on d28, declining together with the advancement of fibrosis (Fig. 5b). Given the different types of sulfation in the total HS composition, a detailed analysis of N-, 2- and 6-sulfation pattern was performed by HPLC (Fig. 5c and Extended Data Fig. 8). Indeed, a marked increase was seen in 6-sulfation (6S) HS portion (Fig. 5c), which is usually associated with a pronounced angiogenic growth^41^. Specifically, 6S HS increased from 2.94±1.11% to 11.87±6.71% (p=0.04) on d7 and to 13.50±5.11% (p=0.012) on d28 post-NSTEMI (Fig. 5c and Extended Data Fig. 8). In addition, a stable significant increase in N-sulfated (NS) HS from 4.82±0.60% to 8.57±1.92% (p=0.035) on d7 and to 11.88±2.63% (p<0.001) on d28 post-NSTEMI (Fig. 5c and Extended Data Fig. 8).

**Figure 5.**
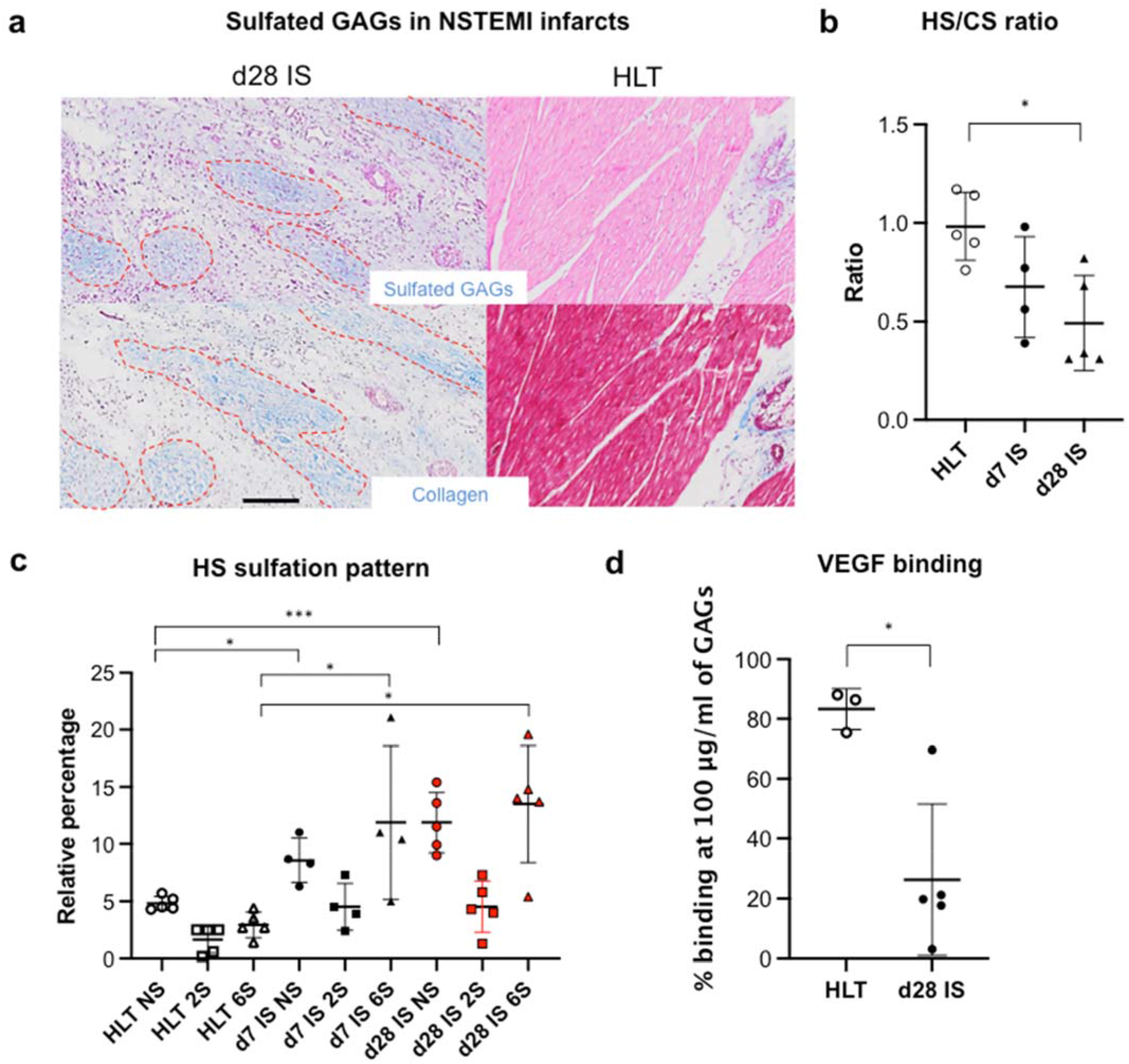
An irreversibly-altered extracellular matrix shows specific changes in the HS sulfation pattern following NSTEMI. **a**, Representative Alcian Blue (top) and Masson’s Trichrome (bottom) stainings to show sulfated glycosaminoglycans (GAGs) and collagen (dashed in red) in ischaemic core (IS) regions of NSTEMI infarcts on d28 post-NSTEMI. *n*=5 animals per group. **b**, Quantification of heparan sulfate (HS) to chondroitin sulfate (CS) ratio in GAGs extracted from tissue. *n*=4 animals at d7 and *n*=5 healthy (HLT) and d28 post-NSTEMI. **c**, Relative percentage of NS, 2S and 6S sulfation in extracted HS across the healthy (HLT) and IS samples at d7 and d28 post-NSTEMI. *n*=5 HLT, *n*=4 at d7 and *n*=5 at d28 post-NSTEMI. **d**, VEGF binding capacity of extracted total GAGs (at 100 µg/ml) across the HLT and IS samples at d28 post-NSTEMI. *n*=3 HLT and *n*=5 animals at d28 post-NSTEMI. Kruskal-Wallis test in (**b**,**c**), Mann-Whitney test in **d**. \**P*<0.05, \*\*\**P*<0.001.

Despite this increase in 6S HS which would suggest angiogenesis, during post-ischaemic remodelling an extended fibrotic replacement compromises physiological vascularity, which is required for normal cardiomyocyte function in healthy conditions. Therefore, to clarify this point in our model of NSTEMI we have looked at the binding capacity of the extracted sulfated HS to angiogenic growth factors, such as VEGF. Binding assays showed a significant drop in the binding capacity 28 days after the surgical procedure (P=0.036) (Fig. 5d), confirming the absence of functional vascularity. In conclusion, ECM GAGs analysis supported the previously observed functional and histological alterations following NSTEMI.

## Discussion

Currently, NSTEMI is the most common presentation of acute MI as most cases with an acute coronary event are NSTEMI patients^4,42-44^. This is partly the result of a widespread use of risk-factor modifying drugs, powerful lipid-lowering statins, and anti-coagulants (aspirin)^45^. NSTEMI patients have lower in-patient (during their admission for the primary NSTEMI) and short-term mortality rates, but significantly higher long-term mortality than those of STEMI patients^6-8^. A Danish registry study of 8.889 patients showed that the 5-year mortality after NSTEMI was 16%^46^, and another registry study highlighted a 10-year survival rate of only around 50%^9^. Nonetheless, to the best of our knowledge, there are currently no large animal models that can reproduce both the functional and histological effects of NSTEMIs as a preclinical base to study interventions that might ameliorate short and long-term effects of NSTEMI. From a preclinical model standpoint, the animal models currently employed usually adopt the ligation of the LAD at different points, and/or including diagonal branches of the LAD and branches of the left circumflex artery which would necessarily produce STEMI and transmural infarcts^10,11^.

In contrast, in the model of NSTEMI presented in this study, multiple ligations (2 cm-apart) lateral and parallel to the LAD were performed from the level of the first diagonal artery - without including it - to within 3-4 cm of the apex to produce patchy and non-transmural infarcts, as evidenced by ECG and histological changes in the antero-lateral wall of the left ventricle. As already well-established in the field, adult sheep were preferred over rodents to achieve a reliable functional outcome effect comparable with that in human cases^20,28^. Indeed, without a relevant preclinical relevant model, interventions using a traditional full-thickness infarction model might lead to clinically ambiguous outcomes^5,47,48^.

The 4th Universal definition of MI requires a rise and fall of cTn and one other criterion from the following: symptoms of acute ischaemia, new ischaemic ECG Changes, new Q-waves, loss of viable myocardium or a new wall motion abnormality in a pattern consistent with ischaemic etiology via imaging^31,49^. In the current model we demonstrated that it is possible to induce infarcts that fulfil all the criteria that define an NSTEMI Type 1 infarct by this multiple suture procedure. Indeed, a significant rise and fall of cTn over time post-MI and the changes typical of NSTEMI on ECG and new wall motion abnormalities on echocardiographic imaging were all present. Moreover, histology showed definitive partial thickness myocardial necrosis and fibrosis. Nonetheless, several important points should be noted in trying to correlate an animal model of NSTEMI with features of NSTEMI seen in the clinics. The absolute peak value of cTn does not consistently correlate with the type of infarction or size of infarction in NSTEMI; no specific cTn level differentiates STEMI from NSTEMI, but cTn values may be used for risk stratification for early intervention^50^. Regarding the evaluation of changes in ECGs, the patterns obtained after the induction of NSTEMI in sheep reflected the typical range of changes classified as NSTEMI in the clinical setting.

Transthoracic echocardiography (TTE) is a non-invasive and well-recognised tool and the protocol employed in this study is based on validated methods^51^. Besides the biological parameters used to determine the response of the infarcted, peri-infarcted and unaffected areas after NSTEMI, functional parameters were also employed to correlate these findings. In addition to EF, which is the most widely used parameter to determine and report left ventricular function, FS was also used. While we acknowledge that the accuracy of FS can be affected by significant apical wall dysfunction or patchy dysfunction as this measurement is taken at one transectional point through the tips of the mitral valve leaflets, which can also sometimes be difficult to pinpoint. FS employs left ventricular end-diastolic and end-systolic diameters (LVEDD and LVESD). These factors are also considered potential markers of left ventricular dilatation in response to myocardial injury. However, LVEDD and LVESD were not primary endpoints since the study endpoint was on day 28 post-NSTEMI, which may not have allowed sufficient time for dilation in response to the myocardial injury. Therefore, regional wall motion index (WMI) analysis was also utilised in this study, and the focus was on the anterolateral walls on TTE. Here we need to consider that scoring is based on a well-validated 3-point scoring system from the American Heart Association^52^, and remain aware that the difference between a normally contracting wall (1), a hypokinetic wall (2) and even an akinetic wall (3) can be subtle in some cases and may vary depending on the observer. We had two different cardiologists review all echocardiograms independently to reduce potential observer error. In case of disagreement there was a discussion and consensus on the scoring.

The current model reflects of clinical presentation in NSTEMI patients who develop myocardial injury and reduction in EF with the significant risk of developing heart failure in the long-term. Specifically, the average decrease in EF observed one (8.52±7.88%) and four (10±8.31%) weeks after the surgical procedure is lower than most of the reductions seen after the total occlusion of the LAD^10,11,27^. The variability in the decrease of this functional parameter reflects the range in clinical cases where multiple factors can influence the functional outcome^22^. A main limitation of this model would be the open thoracotomy nature to create the infarcts. Moreover, although the infarcts are non-transmural, there is a mixture of subendocardial to epicardial infarcts. In particular, epicardial infarcts may impact wall tension differently in the longer-term compared with subendocardial infarcts, resulting in different outcomes in left ventricular geometry.

As previously mentioned, a critical gap in the field of cardiovascular research is the lack of a large animal model that can resemble the clinical cases showing non-full thickness, localised infarcts rather than transmural infarcts in the left ventricular wall. Histological, gene expression and protein analyses indicated that the damaged areas followed the same irreversible fibrotic pattern seen by using STEMI ovine models^20,28^. In addition, extended effects of cardiomyocyte necrosis in the border zone were seen at the mitochondrial level, including the accumulation of dense bodies inside mitochondria, as previously reported after intracoronary balloon occlusion in a swine model^53^.

Post-ischaemic remodelling is tightly coupled with profound alterations in the organisation and composition of the cardiac ECM^33,36^. By proposing a thorough characterisation of the molecular changes occurring in the transcriptome and proteome of the presented NSTEMI model, the relevance of ECM components such as glycoproteins, which were only partially investigated within this context in previous studies^18,54^, has here clearly emerged. Very few studies determined glycosylation alterations post-MI and only following conventional STEMI induction^18,54^. Here, in contrast, using a model of NSTEMI we derived from pathway analyses on gene expression data the relevance of molecular changes in glycans occurring during the post-ischaemic remodelling (day 7 and 28), rather than a pure characterisation of the cardiac post-ischaemic ECM^18,54^. Indeed, technical advances in processing and identifying glycans by mass spectrometry have only recently been developed. These can now be used as tools to elucidate the biological role of glycans^55-57^.

Specifically, LC-ESI-MS/MS on N-linked glycans confirmed the previously reported increase in sialylation^54^, and identified crucial changes such as the marked expression of NeuGc and terminal α-gal only at the inflammatory phase (day 7). Moreover, since most studies evaluating the relevance of NeuGc relate either to cancer biology or to immunotherapy pertaining to xenogeneic reactions^58-60^, data reported in this study constitute a key finding on the expression of NeuGc in myocardial tissue. A relevant study observed a higher NeuGc/NeuAc ratio in adult than in neonatal myocardial tissue^61^. However, this increase in NeuGc was only tenuously associated with cardiomyocyte development, and the precise extent of this change was not defined^61^. In relation to terminal α-gal increase seen in the proposed model on day 7 post-NSTEMI, so far an increased expression of this marker was demonstrated in extensive studies carried out by the Galili group only in wound healing models^62,63^. In addition, α-gal had been found specifically on N-linked glycans in bovine, equine and porcine pericardium^64^. Thus, the increased expression of α-gal in the inflammatory phase post-MI may be mainly due to the temporary recruitment of a disordered and highly proliferative granulation-type tissue. Here, by dissecting the putative N- and O-glycan structures present in the cellular membrane and ECM fractions, we have identified for the first time the precise changes in the glycoprofile pattern through cardiac post-ischaemic remodelling.

Such alterations in glycan expression were also present in ECM components such as GAGs which, unlike the changes in cellular membrane N- and O-glycan composition reported above, contain well-known glycan moieties. Importantly, GAGs are among the structurally fundamental moieties of the ECM; they also play a relevant role in both the TLR-related inflammatory response and the enhancement of angiogenesis through the binding to growth factors^65-67^. We defined how the balance between HS and CS varied post-ischaemia to further advance the current knowledge on cardiac post-ischaemic remodelling. Thus, after quantifying the different GAGs across the remodelling timepoints and by focusing on HS sulfate pattern, we identified increases in markers of angiogenesis on day 7 post-NSTEMI (6S HS). Despite the well-known interaction of 6S HS with angiogenic growth factors^41^, VEGF binding assay data excluded an actual sustained angiogenic effect throughout post-ischaemic remodelling. Therefore, the current model of NSTEMI would include the timely formation of a non-functional vasculature bed in the effected ischaemic area, as it also occurs after STEMI induction in large animals^68^.

This model of non-full thickness MI resembles clinical non-transmural infarcts that are the most prominent type and continue to increase in prevalence among the hospitalised patients. Since there are currently no molecular therapeutic options, the specific alterations in glycans present in the cellular membrane and ECM that we reported at the acute stage post-NSTEMI could be directly targeted by advanced tailored biological interventions to ameliorate the long-term effects of this type of infarction.

## Methods

### NSTEMI ovine model

Regulations established by the European Union directive on the protection of animals for scientific research (2010/63/EU) were followed to perform all the animal experiments presented in the current study. A veterinary team performed all surgical procedures and provided post-operative animal care. To induce NSTEMI, Romanov ten-month-old adult male sheep (35 kg weight on average) were sedated with Telazol^®^ 6 mg/kg (Zoetis, USA) and endotracheally intubated. Anesthesia was maintained with 1–2% isoflurane (Baxter, USA). Electrocardiogram (ECG) was employed to monitor NSTEMI induction throughout the procedure and duration of anaesthesia. Magnesium 2 mg (Pfizer, USA), and amiodarone 1.5 mg/kg (Pfizer, USA) were intravenously injected in sheep before performing the surgical procedure to induce NSTEMI. Following NSTEMI, an intravenous infusion of amiodarone 0.01 mg/kg/min was administered for 1 hour to prevent ventricular arrythmias. During and following surgeries, animals received intramuscular benzylpenicillin 600 mg (Pfizer, USA) three times a day, and streptomycin 500 mg twice daily (Pfizer, USA), for five days. Flunixin meglumine 2.2 mg/kg (Excella GmBH, Germany) was used as analgesic for five days. Whenever lung oedema was detected, hydrocortisone 250 mg intravenous (Pfizer, USA) was administered three times a day.

Induction of NSTEMI was performed by ligating multiple, strategic coronary artery ligations on the LV lateral to and parallel to the LAD. Specifically, a left lateral thoracotomy was performed through the fourth intercostal space, followed by a pericardiotomy. Deep non-transmural ligations were performed with 2/0 Prolene (J&J Ethicon EMEA, Belgium) at 2 cm intervals lateral and parallel to the LAD from the level of the first diagonal moving distally towards and up to 3-4 cm from the apex (see diagram). The blue Prolene sutures used to ligate the coronaries were cut long to allow tracking of the infarction site to aid identification of the sites of infarction. The pericardium was closed with 4/0 Prolene after obtaining absolute hemostasis to limit post-operative adhesions and facilitate re-entry at a later time-point in interventional studies. A chest tube was placed with its tip in the pericardial sac and the remnant holes in the left chest before the thoracotomy was closed in layers, and the animal recovered. The animals were given analgesia and fluids post-operatively as outlined by the institutional protocol. Echocardiography measurements were recorded the day before each surgical procedure and at the study time points of 7 and 28 days. All echocardiographic examinations were performed in calm, unsedated standing animals. A 5 MHz probe was employed and the console and software used were Mindray 7 (Mindray Bio-Medical Electronics Co. Ltd). Images and windows for the echocardiographic protocol were derived from techniques described for horses and more recently adapted for sheep^51^. Two cardiologists performed examinations and agreed on the interpretation and derivation of the data. Six two-dimensional (2-D) parasternal images were obtained from the right, and three 2-D parasternal images from the left. Indices captured were EF, FS, LV volumes and diameters as well as regional wall motion.

### Heart explantation, sampling and tissue processing for histology

At the endpoint of the study (day 28 post-NSTEMI), sheep were anaesthetised as detailed in the description of the surgical induction of NSTEMI. After reopening the thoracotomy wound and pericardium, hearts were explanted. Perfusion with PBS was performed twice to wash out any remaining blood following the explantation. Each explanted heart was sectioned from the ventricular region in an axial way keeping a thickness of 1 cm for each slice. NSTEMI infarcts were visible by whitish colouring of the affected regions in the left ventricle. Explant images were taken with a Cyber-shot DSC-HX200V camera (Sony, Japan). Tissue harvesting was performed by taking multiple samples (0.5 cm maximum) from the ischaemic site, border and remote area. Tissue processing considered the optimal conditions according to the future analysis to be performed. Specifically, samples for histology were submerged in 4% PFA O/N at 4°C. Importantly, histological and immunofluorescence experiments were performed by using a minimum of three sections per sheep, as previously performed^69^.

### Quantification of heparan to chondroitin sulfate ratio

At the time of tissue harvesting around 200 mg of left ventricular tissue from healthy and peri-infarct areas on day 7 and 28 post-NSTEMI were snap-frozen to be processed to extract total sulfated glycosaminoglycans (GAGs). First, dried-powdered samples were weighed and suspended in a buffer to a final concentration of 25 mg of tissue/mL. Tissue digestion was performed by incubating the tissue suspension with proteinase K (PK, 50 μg/mL, Merck, Germany) at 56°C O/N. After enzymatic inactivation at 90°C for 30 min, Dnase (7.5 U/ml, Qiagen, Germany) was added and samples were incubated O/N. Lipid elimination was performed by chloroform extraction, as previously described^40^. After GAGs dialysis, 1,9-dimethylmethylene blue (DMMB) assay was used to quantify GAGs, as already performed^70^. Heparan sulfate (HS) and chondrotin sulfate (CS) quantities were determined by incubating digested samples with a cocktail of heparinases (Iduron, UK), as previously described^71^. Specifically, chondroitinase ABC (25 mU/sample, 2 h at 37 °C) was used for specific CS elimination. Absence of a significant abundance of other GAGs in tissue samples was checked by combining both heparinases and chondroitinases.

### LC-ESI-MS/MS glycomic analysis

After following the procedure to release *N-* and *O-*linked glycans from membrane protein samples, liquid chromatograph-electrospray ionization tandem mass spectrometry (LC-ESI MS/MS) was run to perform the glycomic analysis, as previously reported^72^. A packed in-house column (10 cm × 250 µm) with 5 µm porous graphite particles (Hypercarb™, Thermo Fisher Scientific, USA) was used to separate oligosaccharides. Then, following oligosaccharide injection on to the column, samples were eluted with an ACN gradient (Buffer A, 10 mM ammonium bicarbonate; Buffer B, 10 mM ammonium bicarbonate in 80% ACN). A 40 cm × 50 µm i.d. fused silica capillary was used as a transfer line to the ion source. A Linear Trap Quadrupole (LTQ)-mass spectrometer (Thermo Fisher Scientific, USA), with an IonMax standard ESI source was used to analyse samples in negative ion mode. The heated capillary was kept at 270°C, and the capillary voltage was –50 kV. Each sample was analysed by full scan (m/z 380-2000 two microscans, maximum 100 ms, target value of 30,000), together with MS^2^ scans (two microscans, maximum 100 ms, target value of 10,000) with normalised collision energy of 35%, isolation window of 1.0 units, activation q=0.25 and activation time 30 ms). The threshold for MS^2^ was set to 300 counts. Xcalibur™ software (Version 2.0.7, Thermo Fisher Scientific, USA) was used to perform data acquisition and processing. Identification of the putative glycan structures present in the samples was conducted by manual annotation from their MS/MS spectra. Importantly, assumptions were made to indicate structural annotations. Briefly, N- and O-linked glycan structures were assumed to follow the classic biosynthetic pathways. In addition, diagnostic fragmentation ions to determine N- and O-glycans was performed as previously described^56^. To identify α-linked Gal, terminal hexose-hexose units were considered.

Following MIRAGE guidelines^73^ for glycomic analysis, N- and O-glycan annotated structures were submitted and are currently available at the Unicarb-DB database link https://unicarb-dr.glycosmos.org/references/352. To compare relative N- and O-glycan abundance across different samples, each structure was quantified relative to the total content by integration of the extracted ion chromatogram peak area. Specifically, the area under the curve (AUC) of the N- / O-glycan structure was normalised to the total AUC and indicated as a percentage. Analysis of the peak area was processed by Progenesis QI (Nonlinear Dynamics Ltd, UK).

### nLC-ESI MS/MS label-free proteomic analysis

After digestion and sample preparation, approximately 1.5 μg of peptide per sample was injected into Ultra-High-Performance Liquid Chromatography (UHPLC) system (Ultimate™ 3000 RSLCnano, (Thermo Fisher Scientific, USA) coupled online with Impact HD™ UHR-QqToF (Bruker Daltonics, Germany). Sample analysis was performed twice to reduce variability. Sample loading was conducted by using first a pre-column (Dionex, Acclaim PepMap 100 C18, cartridge, 300 µm), and then a 50 cm nano-column (Dionex, ID 0.075mm, Acclaim PepMap100, C18). Sample separation occurred at 40°C with a flow rate of 300 nL/min using multistep 4h gradients of ACN, as already reported^74^. The column was connected to a nanoBoosterCaptiveSpray™ ESI source (Bruker Daltonics, Germany). Collision-induced Dissociation (CID) fragmentation (N2 as collision gas) was applied as data-dependent acquisition mode. Before each sample was run, a specific lock mass (1221.9906 m/z) and a calibration segment (10 mM sodium formate cluster solution) were applied to improve mass accuracy. Data was acquired as previously described^75^.

DataAnalysis™ v.4.1 Sp4 (Bruker Daltonics, Germany) was used to elaborate data and protein identities and relative abundancies were determined using Peaks Studio 8.5 (Bioinformatics Solutions Inc., USA)^76^. Each sample was run and analysed as two independent replicates. For protein identification, Uniprot’s reference database of *Ovis aries* was accessed on Feb 2018, 556,825 sequences; 199,652,254 residues. In particular, the following parameters were set: enzymatic digestion performed by trypsin, allowing one missed cleavage; precursor mass tolerance was 20 ppm; fragment mass tolerance of 0.05 Da; carbamidomethylation as fixed modification. False-positive identification rate (FDR) was set ≤1%, and a peptide score of −log10(p-value) ≥ 20 was considered adequate for confident protein identification. De Novo ALC score was set ≥ 50%. Relative peptide signal intensity was calculated only for confidently identified peptide features. Then, AUC of each extracted ion chromatogram was calculated and used for the relative quantification after normalisation Total Ion Current (TIC). Cumulative peak areas of the proteins were measured by considering only unique peptides assigned to specific proteins. Only proteins with more than two unique peptides were considered for the analysis.

### Statistical analysis

The *in vivo* study was designed and powered for statistical significance (power = 0.8), and this required a minimum of six animals per group. Animals from at least two different batches of surgeries were analysed for each group in all analyses to avoid a single batch-dependent bias. Data are presented either as box plots or as the average ± standard deviation (s.d.) and differences with *P*<0.05 were considered significant. Statistical analyses were performed using Prism^®^ v9.0.0 (GraphPad Software Inc., USA). Normality and equality of variance were tested before a statistical test and either Student’s *t* - test or Mann-Whitney *U* - test accordingly used when comparing two groups. Multiple group analyses were performed by running either Kruskal-Wallis or Tukey’s one-way ANOVA after assessing normality and equality. For RNA-seq analysis, the Wald test was used to generate p-values and Log2 fold changes as per DEseq2 method. Statistical differences were defined as \**P*≤0.05, \*\**P*≤0.01, \*\*\**P*≤0.001.

## Data availability

The authors declare that all data supporting the findings of this study are available from the authors on reasonable request. RNA-seq raw data are publicly available in GEO under accession code GSE164245. Glycomics-derived annotated structures were deposited in the Unicarb-DB database (https://unicarb-dr.glycosmos.org/references/352).

## Code availability statement

No custom computer code or mathematical algorithm that is deemed central to the conclusions was used in this study.

## Acknowledgements

The authors acknowledge the use of the Centre for Microscopy and Imaging facilities at the National University of Ireland Galway. Mass Spectroscopy analysis of glycans was performed by the Swedish infrastructure for biological mass spectrometry (BioMS) supported by the Swedish Research Council. P. Lalor and E. McDermott from the Anatomy at the National University of Ireland Galway assisted in processing for TEM analysis. R. Grigalevičiūtė and A. Kučinskas from the Lithuanian University of Health Sciences assisted with veterinary procedures and in ensuring animal well-being. Prof. D.H. Pauza and Dr K. Rysevaite from the Institute of Anatomy at the Lithuanian University of Health Sciences (Kaunas, Lithuania) assisted with initial processing of tissue samples. The authors would like to acknowledge A. Sloan and Dr R. Bohara for editorial assistance, M. Doczyk for assistance in drawing the schematics presented in this article and Dr O. Carroll for technical help. This manuscript is dedicated to Mimì.

## Funding

This work was funded by the European Commission funding under the AngioMatTrain 7^th^ Framework Programme (Grant Agreement Number 317304) and by the research grant from Science Foundation Ireland (SFI) co-funded under the European Regional Development Fund under Grant Number 13/RC/2073 and13/RC/2073_P2.

## Author contributions

P.C., M.D.C. and A.P. conceived the idea and designed the experiments. M.D.C., V.V., M.R. and P.C. developed and validated the ovine model of MI and performed the *in vivo* study. P.C. performed the *in vivo* experiments and relative analyses. A.K. and E.E. assisted in functional data recording and analysis. V.Z. coordinated all veterinary procedures and animal welfare. A.G.P. assisted with the *in vivo* experiments. P.C. and R.S. ran analyses on gene expression and proteomic data. C.J. and N.G.K. performed the PG-LC-ESI-MS/MS glycomic analysis. C.C. and F.M. performed the nLC-ESI-MS/MS proteomic analysis. S.C. and D.P-G. performed the GAGs analysis. P.C. wrote the manuscript. M.D.C. edited the manuscript. A.P. supervised the entire project. All authors read the manuscript, commented on it and approved its content.

## Declaration of Competing Interest

The authors declare no competing financial interests or personal relationships that could have appeared to influence the work reported in this paper.

## Notes

### Competing Interest Statement

The authors have declared no competing interest.

