## Supplementary material for "Reproducing extracellular matrix adverse remodelling of non-ST myocardial infarction in a large animal model": Supplimentary figure 1

### Supplementary Information

#### Supplementary Methods

##### Electrocardiograms data from NSTEMI patients

The electrocardiograms (ECGs) which validated functional changes consistent with induced NSTEMI in sheep (Extended Data Fig. 2b) were standard 12-lead ECGs in human subjects. These were patients who presented with signs and symptoms suggestive of an MI which were subsequently confirmed on the ECG and serum troponin measurements at University Hospital Galway (Ireland). All personal data identifiers have been removed for obvious reasons.

The protocol for obtaining consistent ECGs is based on hospital protocols derived from AHA recommendations[1-3]. Many patients are uncomfortable lying flat, so for consistency and practicality, a semi-recumbent position of approximately 45 degrees is recommended. There is no evidence that variation of the inclination of the patient between horizontal and 45 degrees to the horizontal has any significant effect on the ECG. Patients with excessive hair in small areas corresponding to where leads had to be placed had these areas shaved to ensure optimal adhesion and contact of the electrodes. The skin was cleansed with mild soap and water or alcohol swabs to remove oil that may be on the skin and which can cause drift in the ECG signals. Adhesive electrodes were then attached to the patients in the standard positions: limb leads on the right arm, left arm, right leg and left leg and the precordial or ventricular leads V1-V6 at the standard positions across the chest. All 12-lead ECGs and simultaneous rhythm strips were recorded at 25 mm/s with a gain setting of 10 mm/mV. The patients were instructed to breathe normally and not to talk or move during the short period the ECG was being captured to avoid muscle activity causing motion artifact.

**Electron microscopy analysis**

On d7 and d28 post-NSTEMI induction tissue samples from healthy and ischaemic, border and remote infarcted hearts were fixed in 2.5% gluteraldehyde at 4°C for 24h. Tissue processing for transmission electron microscopy analysis (TEM) started by washing samples in phosphate buffer (PB) and post-fixing in 1% osmium tetraoxide (Sigma Aldrich, USA) for 2 h at RT. Following this, samples were gradually dehydrated in 30%, 50%, 70%, 95% and pure ethanol. Then, two washes in pure acetone were performed before the embedding. Specifically, samples to be embedded were first infiltrated with Araldite® epoxy resin (Electron Microscopy Sciences, USA) at a progressive 2:1, 1:2 ratio with acetone and finally with pure resin for 24 h each at RT. To conclude the processing, samples were moved to freshly made pure Araldite® epoxy resin, then samples were left to cross-link at 60°C for 48 h. Once embedded, ultrathin sections (70-90 nm thick) were cut into samples using a diamond glass knife and were then put onto copper grids. Tissue sections were post-stained with 0.5% uranyl acetate (Laboratory Instruments & Supplies, Ireland) for 3 min and 3% lead citrate (Laboratory Instruments & Supplies, Ireland) for 5 min using an EM AC20 Auto Ultrastainer® (Leica, Germany), as previously performed[4].

**Release of N- and O-linked glycans from membrane proteins**

Snap-frozen healthy and infarcted (ischaemic core, border and remote regions) were digested using the Mem-PER™ Plus Membrane Protein Extraction Kit (Thermo Fisher Scientific, USA) to extract cytosolic and membrane protein fractions as per manufacturer's instructions. Protein concentration was determined using a Micro BCA™ Protein Assay Kit (Thermo Fisher Scientific, USA). To extract N-linked glycans from membrane protein fractions, ischaemic, border and remote region samples derived from three different animals per group were used. Protein extraction buffer was exchanged to 7 M urea through a 30 kDa MWCO centrifugal filter (Millipore, USA), followed by an incubation with 25 mM dithiothreitol (DTT) at 56°C for 45 min. After reduction, samples were alkylated with 62.5 mM iodoacetamide (IAA) at RT for 50 min in the dark. Then, samples were trypsinised (sequencing grade 1% w/w, Promega, USA) O/N at 37°C and resulting peptides were precipitated with 80% (v/v) acetone. Pellets were left to dry, then washed twice with cold 60% methanol and again air-dried. To release N-linked glycans, samples were incubated with PNGase F (Asparia Glycomics, Spain) in ammonium acetate (50 mM, pH 8.4) O/N at 37°C. Then, a SEP-Pak C18 cartridge (Waters Corporation, USA) was used to separate N-linked glycans from (*O*-glyco)peptides. Specifically, a SEP-Pak C18 cartridge was conditioned with dilutions (90% and 10%) of acetonitrile (ACN) in 0.5% trifluoroacetic acid (TFA). Once the sample was applied, elution with 5% acetic acid allowed the release of N-linked glycans. Moreover, elution of O-glycopeptides was conducted by adding 65% ACN in 0.5% TFA. After drying at 45°C, N-linked glycans were reduced by 0.5 M sodium borohydride ($NaBH_4$) and 20 mM NaOH O/N at 50°C. In addition, reductive β-elimination reaction enabled the release of O-linked glycans by incubation in a buffer containing 0.5 M $NaBH_4$ and 50 mM NaOH O/N at 50°C. Reaction quenching was performed with glacial acetic acid. Finally, samples were desalted and dried as previously described[5].

**Protein extraction and digestion for proteomic analysis**

Ischaemic, border and remote region samples (30-50 mg) from NSTEMI-infarcted sheep were snap-frozen following heart explantation. On the day of tissue processing, samples were thawed and finely cut in small pieces with a scalpel before starting tissue digestion and protein extraction. RIPA lysis and extraction buffer (Thermo Fisher Scientific, USA) with cOmplete™ EDTA-free protease inhibitor cocktail (Roche, Switzerland) was used to resuspend the samples before homogenisation (2 cycles 15 min each) using a TissueLyser LT (Qiagen, Germany) set at 50 oscillations per min. Once completely homogenised, samples were incubated on ice for 15 min and centrifuged at 12,000 rpm for 10 min at 4°C. Total protein concentration was determined by Micro BCA™ Protein Assay Kit (Thermo Fisher Scientific, USA) after setting a titration curve. Then, protein samples were concentrated and detergents removed by a suspension trapping (S-Trap) method[6]. Specifically, S-trap™ Micro spin columns (ProtiFi, USA) were used following manufacturer's instructions, with minor modifications. Briefly, Tris-HCl was used instead of TEAB (Triethylammonium bicarbonate) buffer. Approximately 100 ug of protein sample was dried and resuspended for further reduction (20 mM DTT, 10 min 95°C) and alkylation (40 mM IAA, 30 min in the dark). After incubation in aqueous phosphoric acid at 1.2% final concentration, binding buffer was added and samples were loaded into micro-columns for protein trapping and trypsinisation (1h at 37°C), as per manufacturer's protocol. Peptides elution was performed by centrifuging at 4,000 g with ammonium bicarbonate (50 mM) and 0,2% formic acid (FA). Recovery of hydophobic peptides was performed with 50% ACN with 0.2% FA. Final peptide concentration was determined using a NanoDrop™ One/Onec Microvolume UV-Vis Spectrophotometer (Thermo Fisher Scientific, USA).

**RNA sequencing**

Following tissue harvesting, samples (0.5 cm maximum) from the ischaemic site, border and remote areas of infarcted or from left ventricular wall healthy hearts were harvested and stored in RNA*later*™ (Thermo Fisher Scientific, USA) at -80°C until further processing. Each tissue sample (40 mg) was finely cut with a scalpel, digested in TRIzol™ (Thermo Fisher Scientific, USA) by bead grinding using a TissueLyser LT (Qiagen, Germany) and extracted by the phenol-chloroform method. Specifically, once samples were completely homogenised, RNA was extracted using RNeasy® Mini Spin Columns (Qiagen, Germany), as per manufacturer's instructions.

RNA sample yield, quality and integrity were assessed by Qubit® 2.0 Fluorometer (Life Technologies, USA) and Agilent TapeStation (Agilent Technologies, USA). RNA library preparation, sequencing and bioinformatics analysis were conducted at GENEWIZ, Inc. (USA). Briefly, NEBNext® Ultra™ RNA Library Prep Kit for Illumina was used for RNA-seq library preparation. Agilent TapeStation (Agilent Technologies, USA) was employed to validate the sequency library and cDNA were quantified by using Qubit® 2.0 Fluorometer (Invitrogen, USA) as well as by quantitative PCR (KAPA Biosystems, USA). Sequencing libraries were loaded on the Illumina® HiSeq 4000 in high output mode and samples sequenced using a 2x150 paired end configuration. Image analysis and base calling were conducted by the HiSeq Control Software. Raw sequence data (.bcl files) generated from Illumina® HiSeq was converted into fastq files and de-multiplexed using Illumina's bcl2fastq 2.17 software. One mis-match was allowed for index sequence identification. Sequence reads were trimmed to exclude adapter sequences and poor quality nucleotides using Trimmomatic v.0.36. Reads mapping was performed using STAR aligner v.2.5.2b on the human reference genome available on ENSEMBL. Unique gene hit counts were calculated by using feature Counts from the Subread package v.1.5.2 and counted only when falling within exon regions. Following gene hit count extraction, data was used to perform differential expression gene (DEG) analysis. Comparison of gene expression between the groups of samples was performed within the package DESeq2.

**Ingenuity Pathway Analysis (IPA®) analysis of transcriptomic and proteomic data**

DEG data were processed as .xls files. The gene list was sorted to set all the data relative to the different groups to a summed-up ID gene list. The created .xls file had columns reporting the log2(fold change) DEG data of each condition compared to the healthy animals. This file was loaded in IPA® (Qiagen, Germany) software, as per manufacturer's instructions. Log2(fold change) > 1.5 and < -1.5 and adjusted P-value < 0.05 were applied as cut-offs for core analysis. Comparative analysis was conducted on the files of core ischaemic samples on days 7 and 28 post-NSTEMI. Canonical pathway and upstream regulators data were compared by their activation z-score and associated p-value. Identified protein data from nLC-ESI-MS/MS label-free quantification with a unique peptide number above 2 were selected and listed in a .xls file. Protein entries from the infarcted samples were reported in terms of fold change ratio to healthy conditions. The .xls file was loaded in the IPA® (Qiagen, Germany) software according to the instructions. Cut-offs of Fold change > 1.5 and < -1.5 were applied for core analysis.

#### Supplementary Figures and Table

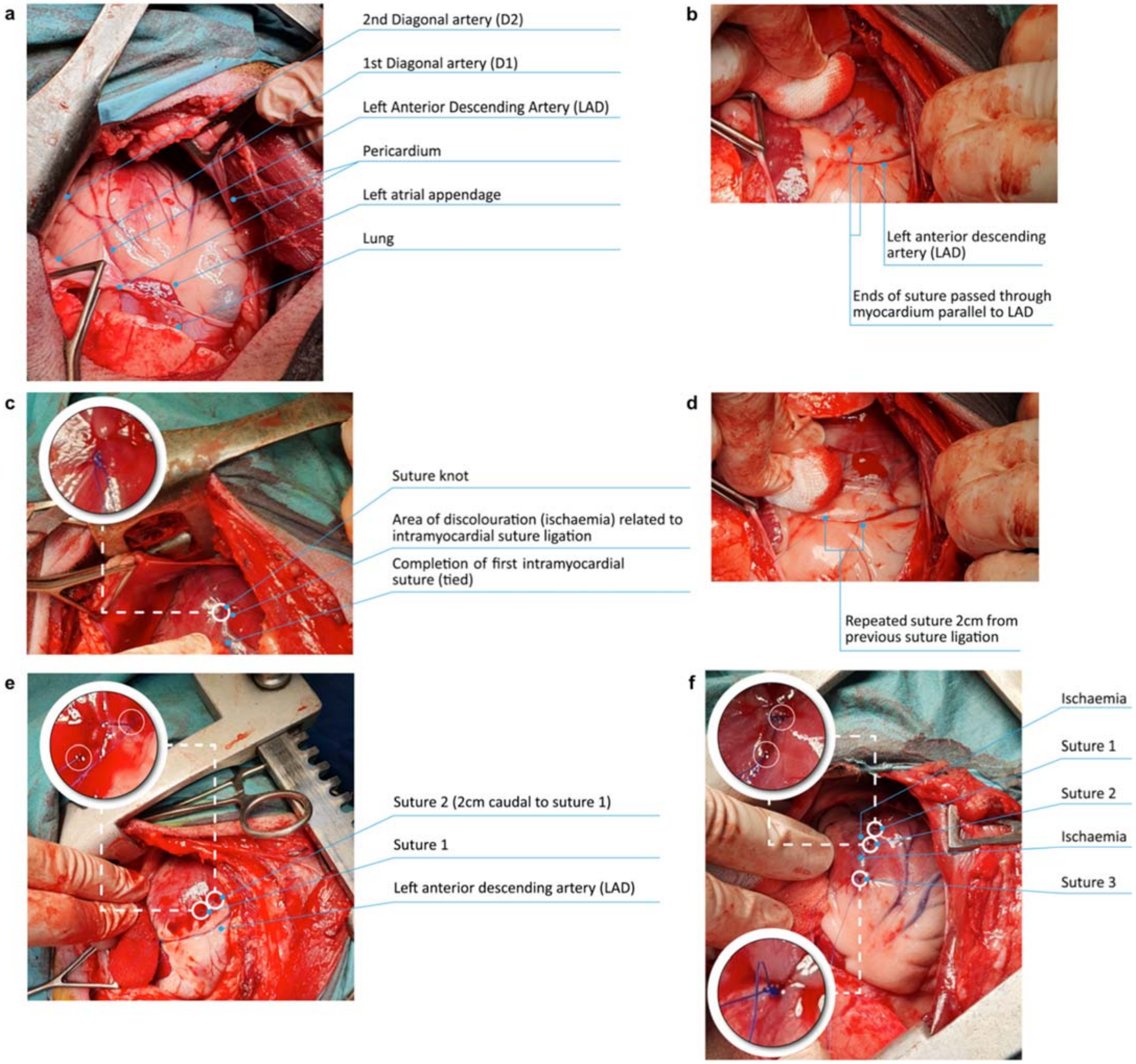

**Extended Data Figure 1 | Surgical procedure of NSTEMI induction in sheep**

**a**, Following thoracotomy and pericardiectomy, the heart is exposed to detect LAD and diagonal arteries. **b**, A suture is passed through the left ventricular myocardium parallel to the LAD. **c**, The first suture knot is completed and an initial area of discoloration appears in the proximity of the ligated ventricular portion. **d-e**, A second suture is performed approximately 2 cm apart from the first one, always parallel to LAD. **f**, Progressively, a third suture is performed and tied in the same direction towards the apex of the heart again 2cm apart from the previous suture. The pale area of ischemia across the suture knots is clearly evident.

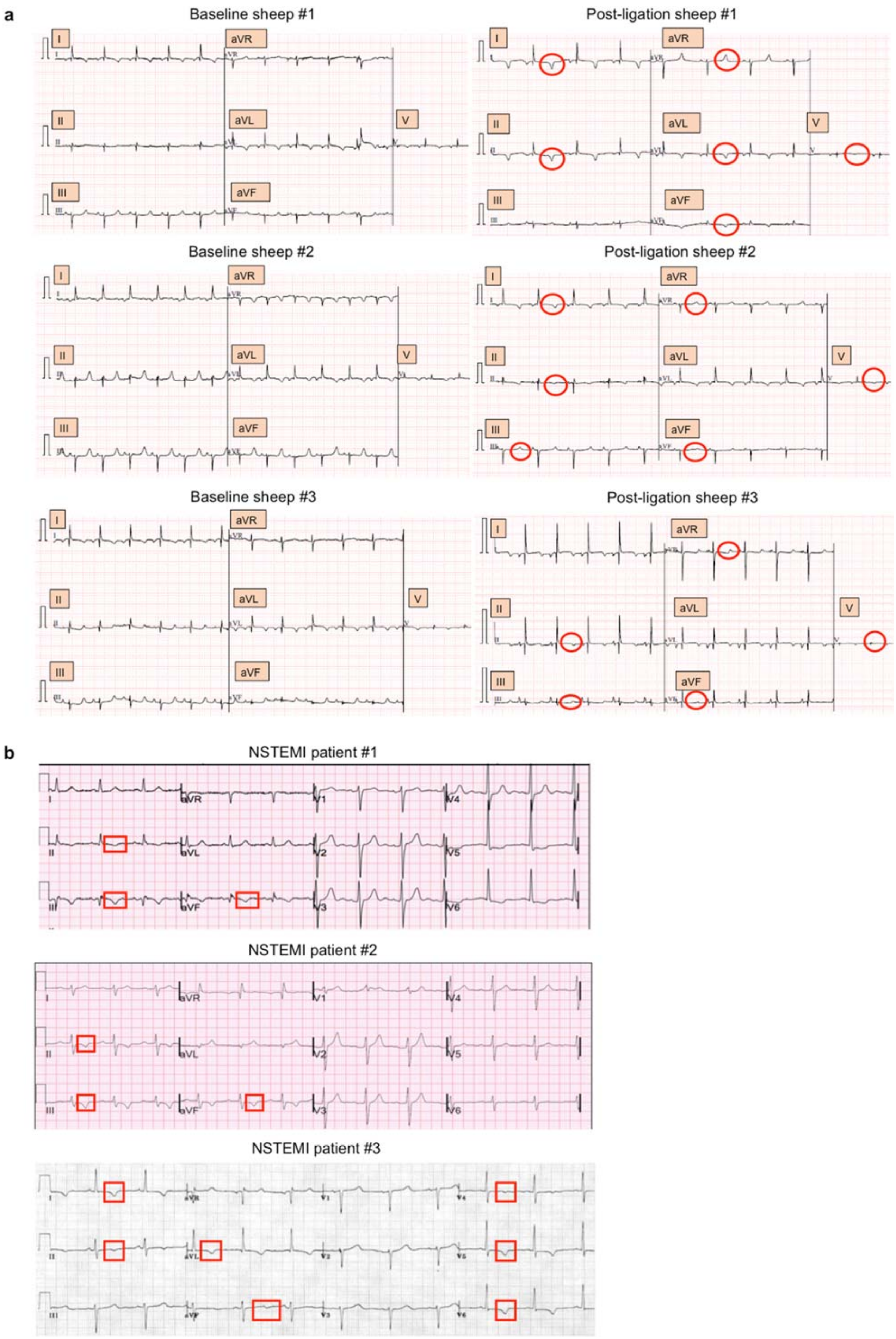
a
Baseline sheep #1
Post-ligation sheep #1
Baseline sheep #2
Post-ligation sheep #2
Baseline sheep #3
Post-ligation sheep #3
I
aVR
II
aVL
V
III
aVF
b
NSTEMI patient #1
NSTEMI patient #2
NSTEMI patient #3

**Extended Data Figure 2 | Resemblance of NSTEMI model to clinical cases**

**a**, ECG tracings before and after the induction of an NSTEMI type infarction in sheep. Baseline: ECG tracing with no significant changes in QRS, ST segments or T waves. Post-ligation changes (circled in red) in sheep #1: T wave inversion in leads II and aVF, accentuated peaking of T waves in I and aVR, with flattening of the T wave in leads III and V. Post-ligation changes (circled in red) in sheep #2: T wave inversion in leads I, II and eversion of T wave in aVR, with flattening of the T waves in leads III, aVF and V. Post-ligation changes (circled in red) in sheep #3: T wave inversion in leads II, III and eversion of T wave in aVR, with flattening of the T waves in leads II, aVF and V. **b**, Clinical ECG tracings in three different patients diagnosed with NSTEMI. Top, patient #1, 59 year old complaining of persistent angina for two hours: T wave inversion with ST depression (framed in red) in leads II, III and aVF. Middle, patient #2, 67 year old presenting with prolonged chest pain of sudden onset: T wave inversion with ST depression (framed in red) in the inferior leads II, III and aVF. Bottom, patient #3, 72 year old with acute onset of prolonged chest pain: Widespread T wave inversion (framed in red) in the septal and inferolateral leads.

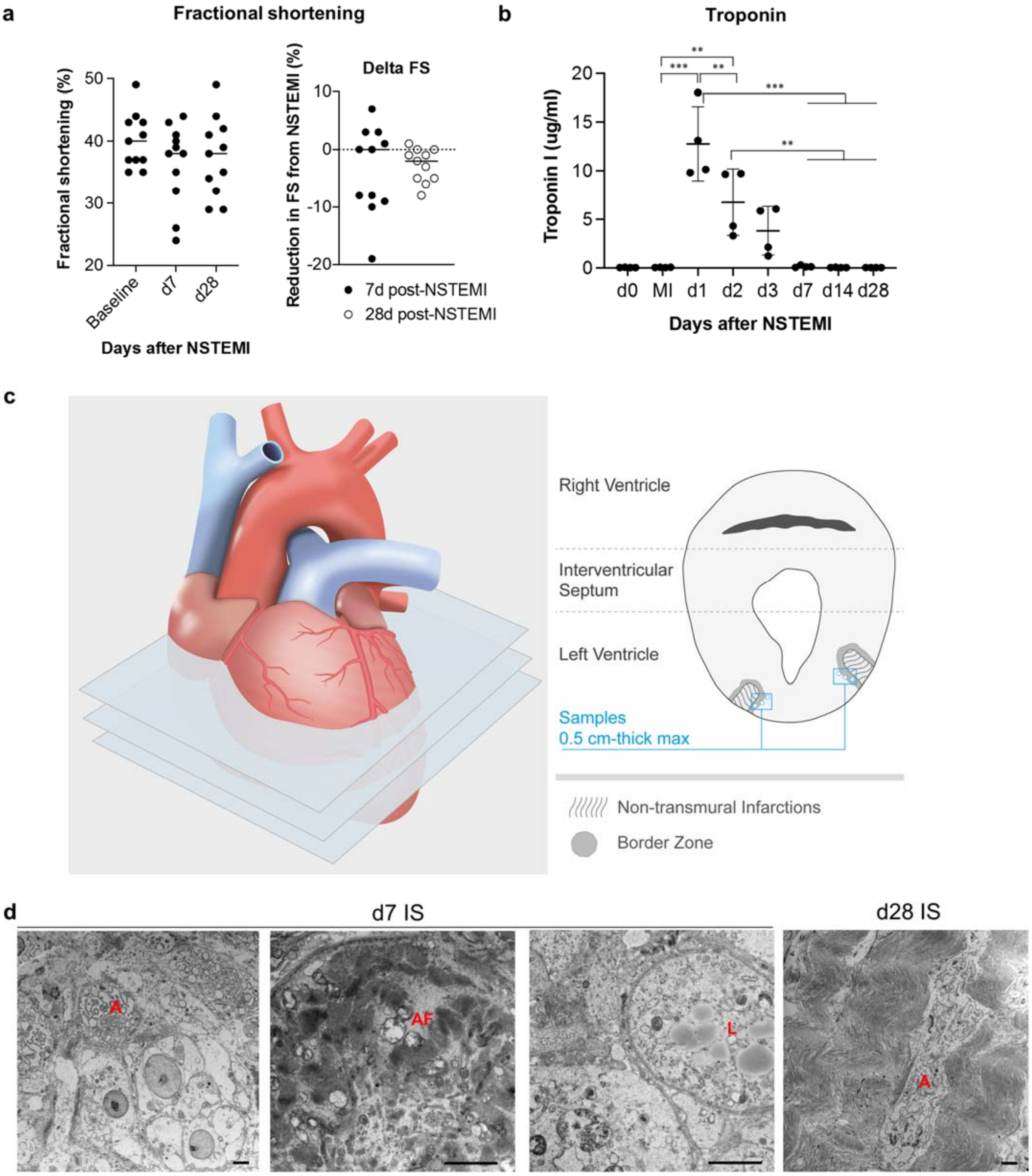

**Extended Data Figure 3 | Functional and histological evaluation of NSTEMI model**

**a**, Fractional shortening (FS) percentage at baseline, 7 and 28 days post-NSTEMI. Reduction in FS (delta) at day 7 and 28 after surgery. *n*=11 animals. **b**, Troponin measurements before surgery and throughout the four weeks following NSTEMI induction. *n*=4 animals. **c**, Schematics of tissue harvesting from explanted hearts. After perfusion with PBS to remove the excess of blood, each heart was cross-sectioned in slices with a thickness of 1 cm from the atria to the apex. Clear MI areas were identified by the whitish colour and from there samples with a maximum size of 0.5 cm in every direction were taken from the core ischaemic, the border and the remote regions. **d**, Representative TEM micrographs at 7 days post-NSTEMI: Apoptotic bodies (A), autophagosomes (AF) and lipid droplets (L) are widely spread in ischaemic regions previously populated by intact cardiomyocytes. Representative TEM micrographs at 28 days post-NSTEMI: Wavy-oriented typical collagen-like deposition replaced cardiomyocytes. *n*=5 animals per group, scale bar = 2 μm. Data in **a-b** are reported as dot-plots, one-way ANOVA with Tukey's post hoc correction, Wilcoxon test for delta FS. **p<0.01, ***p<0.001.

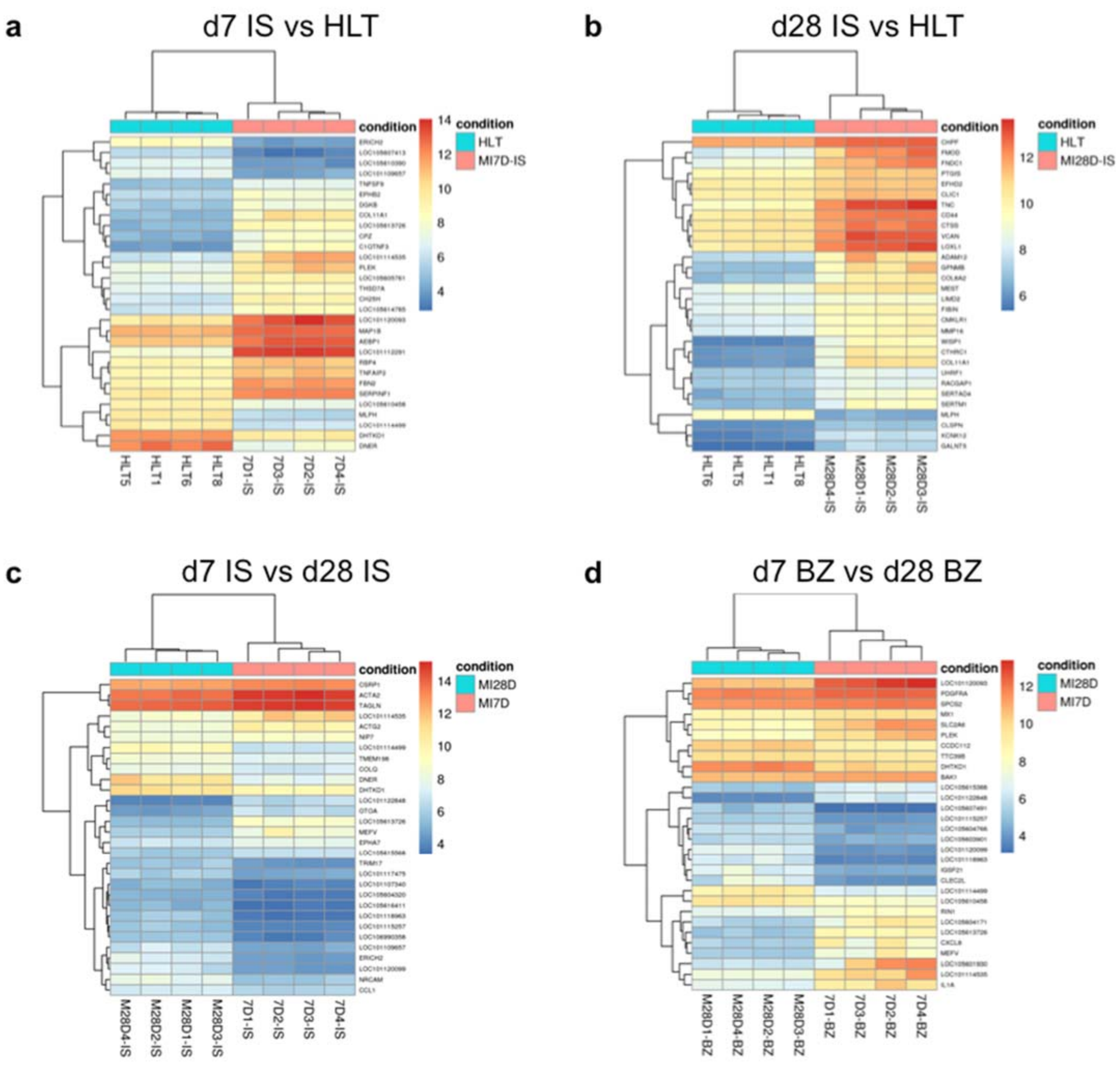

**Extended Data Figure 4 | Transcriptomics in ischaemic core following NSTEMI**

**a**,**b** Bi-clustering heat map listing the top 30 genes that are called differentially expressed (DEG) sorted by lowest p-values ($p<0.05$) detected in ischaemic core at day 7 (**a**) and 28 (**b**) post-NSTEMI compared with healthy (HLT) myocardial ventricular tissue from whole RNA-sequencing. **c**,**d** Bi-clustering heat map of DEG between 7 and 28 days post-NSTEMI in the ischaemic core (**c**) and in the border zone (**d**). Difference in mRNA expression between the conditions is indicated as log2 (fold change). Analysis was performed using DESeq2 package and statistical significance assessed by Wald's test. Adjusted-$p<0.05$ was set to identify differences. *n*=4 animals per group.

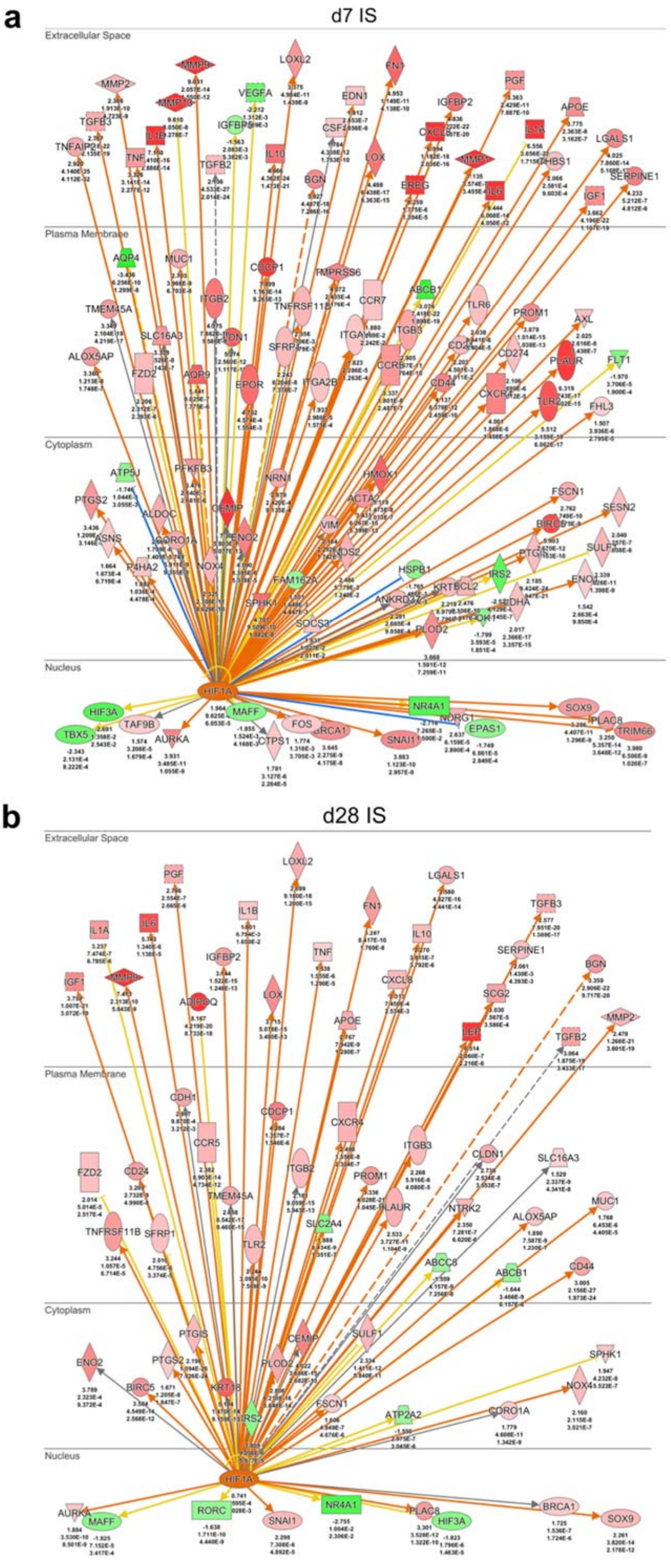

a
d7 IS
Extracellular Space
MMP2
MMP9
MMP13
TGFB3
IL1B
TNF
VEGFA
TGFB2
LOXL2
IL10
EDN1
BGN
LOX
FN1
IGFBP2
EREG
IL6
PGF
IL1A
THBS1
APOE
LGALS1
SERPINE1
IGF1
Plasma Membrane
AQP4
MUC1
TMEM45A
ITGB2
CDCP1
TMPRSS6
CCR7
ABCB1
TLR6
PROM1
AXL
SLC16A3
ALOX5AP
FZD2
AQP9
EPOR
ITGA2B
ITGB3
CD44
CD274
PLAUR
FLT1
FHL3
Cytoplasm
ATP5J
PFKFB3
NRN1
HMOX1
ACTA2
FSCN1
SESN2
PTGS2
ALDOC
CEMIP
ENO2
VIM
BIRC5
ASNS
CORO1A
P4HA2
NOX4
HSPB1
IRS2
SPHK1
FAM162A
SOCS3
PLOD2
Nucleus
HIF1A
HIF3A
TAF9B
MAFF
NR4A1
SOX9
TBX5
AURKA
CTPS1
FOS
SNAI1
EPAS1
PLAC8
TRIM66
b
d28 IS
Extracellular Space
PGF
LOXL2
LGALS1
IL1A
IL6
IL1B
FN1
TGFB3
IL10
IGFBP2
TNF
SERPINE1
IGF1
MMP9
LOX
CXCL8
BGN
ADIPOQ
SCG2
APOE
LEP
MMP2
TGFB2
Plasma Membrane
CDH1
CDCP1
CXCR4
CCR5
ITGB3
CLDN1
SLC16A3
FZD2
CD24
ITGB2
PROM1
TMEM45A
PLAUR
NTRK2
SLC2A4
ALOX5AP
MUC1
TNFRSF11B
SFRP1
TLR2
ABCC8
ABCB1
CD44
Cytoplasm
PTGIS
CEMIP
SULF1
SPHK1
ENO2
PTGS2
PLOD2
BIRC5
KRT18
IRS2
FSCN1
ATP2A2
CORO1A
NOX4
Nucleus
HIF1A
AURKA
MAFF
RORC
SNAI1
NR4A1
PLAC8
HIF3A
BRCA1
SOX9

**Extended Data Figure 5 | HIF1□ pathway analysis in ischaemic core following NSTEMI**

**a**,**b** HIF1$\alpha$ is a key regulator in NSTEMI at day 7 (**a**) and 28 (**b**) days post-surgery. Shades of red and green indicate the degree of upregulation and downregulation, respectively. The activation of the downstream node is orange-coded, inhibition is blue-coded: if the findings underlying the relationship are inconsistent with the state of the downstream node yellow is used, and grey if there is no predicted effect. Pointed arrowheads indicate that the downstream node is expected to be activated, while blunt arrowheads indicate that it is expected to be inhibited. Log2(fold change), P, and adjusted-P from RNA-seq data are indicated below the gene names. *n*=4 animals per group.

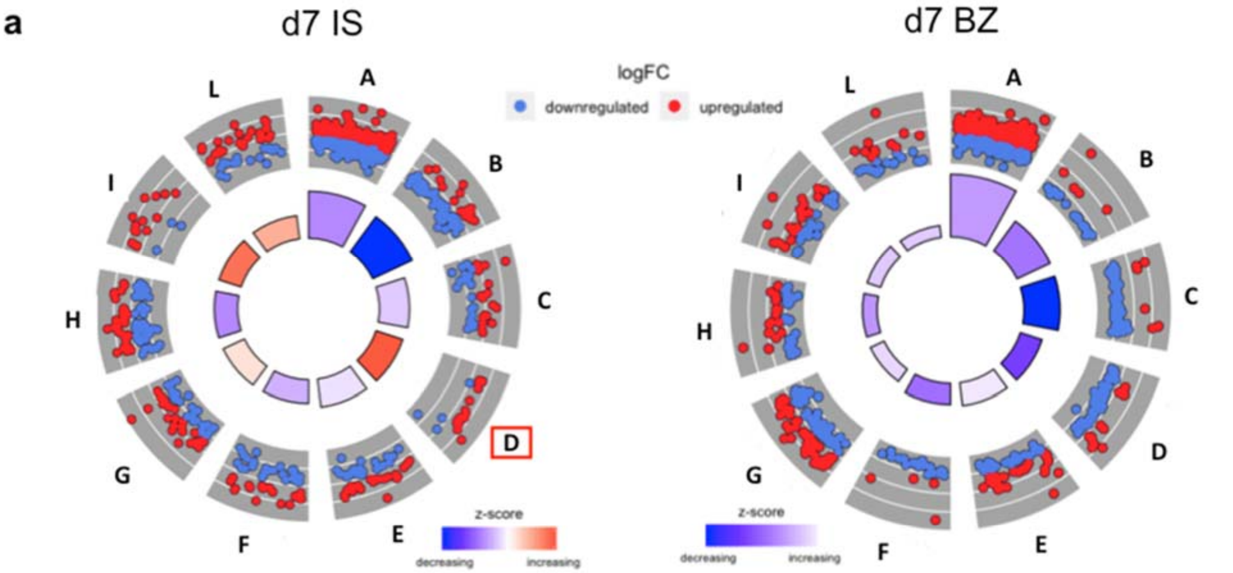

| ID | Description |
|---|---|
| A | Metabolic pathways |
| B | Ubiquitin mediated proteolysis |
| C | Glycerophospholipid metabolism |
| D | N-Glycan biosynthesis |
| E | Neurotrophin signaling pathway |
| F | FoxO signaling pathway |
| G | Hippo signaling pathway |
| H | MAPK signaling pathway |
| I | DNA replication |
| L | Cell cycle |

| ID | Description |
|---|---|
| A | Metabolic pathways |
| B | Citrate cycle (TCA cycle) |
| C | Oxidative phosphorylation |
| D | Ubiquitin mediated proteolysis |
| E | FoxO signaling pathway |
| F | Pyruvate metabolism |
| G | MAPK signaling pathway |
| H | Insulin signaling pathway |
| I | AMPK signaling pathway |
| L | Glycolysis / Gluconeogenesis |

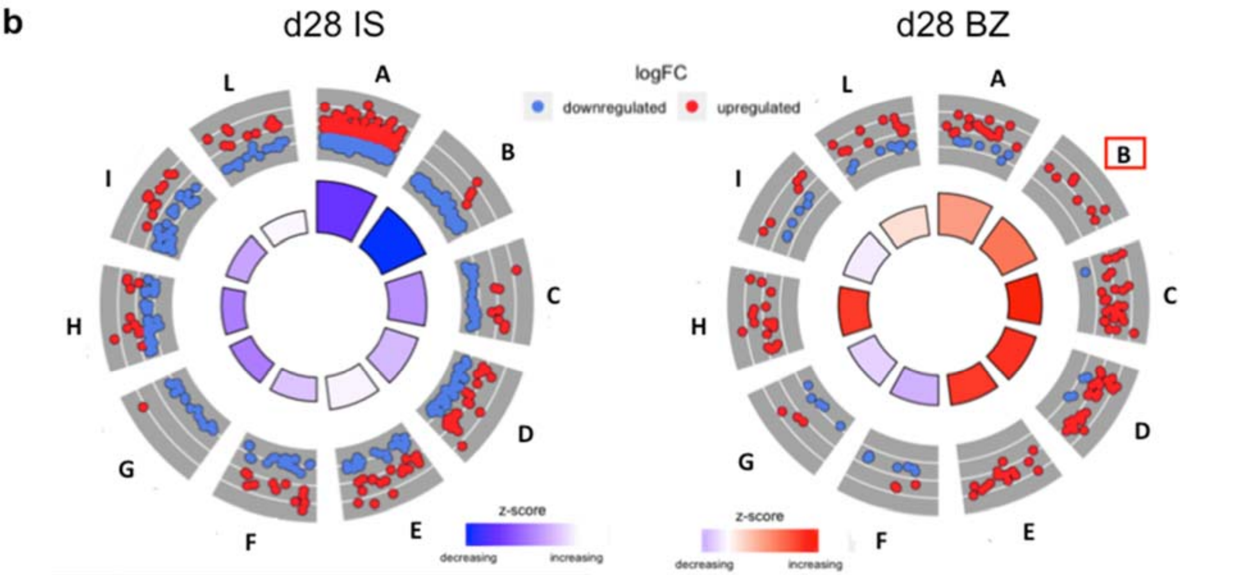

| ID | Description |
|---|---|
| A | Metabolic pathways |
| B | Oxidative phosphorylation |
| C | Cardiac muscle contraction |
| D | Adrenergic signaling in cardiomyocytes |
| E | Glycerophospholipid metabolism |
| F | Dilated cardiomyopathy |
| G | Citrate cycle (TCA cycle) |
| H | Insulin signaling pathway |
| I | Glucagon signaling pathway |
| L | Arrhythmogenic RV cardiomyopathy |

| ID | Description |
|---|---|
| A | Cell cycle |
| B | Glycosaminoglycan biosynthesis |
| C | Chemokine signaling pathway |
| D | Focal adhesion |
| E | Fc gamma R-mediated phagocytosis |
| F | Fatty acid degradation |
| G | Proponoate metabolism |
| H | ECM-receptor interaction |
| I | Fatty acid metabolism |
| L | FoxO signaling pathway |

**Extended Data Figure 6 | Functional annotation analysis from RNA-seq of NSTEMI model**

**a**,**b** Functional annotation analysis using DAVID software of the DEG in ischaemic core and border zone following NSTEMI. Gene-annotation enrichment analysis (KEGG_PATHWAY) of the set of DEG (adjusted p-value<0.05) is displayed using the GOCircle plot in which the inner ring is a bar plot where the height of the bar indicates the significance of the term (log10 adjusted p-value), and the colour corresponds to the z-score, calculated as the number of up-regulated genes minus the number of down-regulated genes divided by the square root of the upregulated and downregulated genes; red dots represent up-regulated genes and blue dots indicate down-regulated genes. **a**, N-glycan biosynthesis (framed in red) shows the highest z-score in the ischaemic core at day 7 post-surgery. **b**, Glycosaminoglycan biosynthesis (framed in red) is one of the top biological cathegories showing a highly activated z-score in the border zone at day 28 post-surgery. n=4 animals per group.

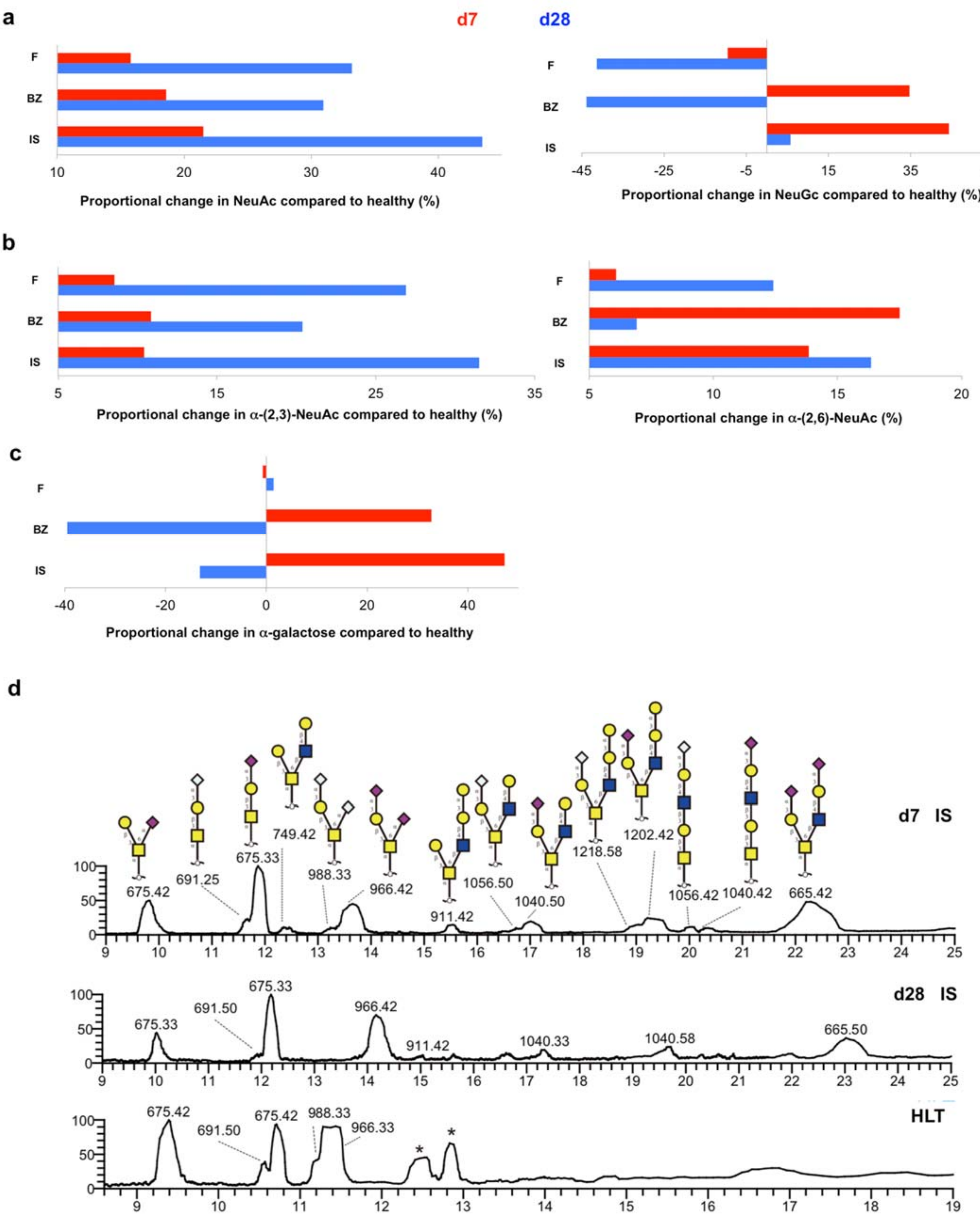

**Extended Data Figure 7 | Glycomics analysis of infarcted hearts following NSTEMI**

Proportional changes in N-linked glycans in the ischaemic core, border and remote regions compared to healthy. **a**,**b** Amount of N-linked glycans expressing either NeuAc or NeuGc sialic acid (**a**) and linkage type (**b**) α(2,3) or α(2,6) across the different regions following NSTEMI at day 7 and 28 post-surgery. **c**,

Amount of terminal α-gal in the same infarcted regions. **d**, Extracted ion chromatography (EIC) showing O-linked glycans mainly expressed in the membrane protein extracts from the ischaemic core at days 7 and 28 post-surgery compared with healthy (HLT) myocardial tissue. Regions of infarcted hearts are labelled as follows: IS = core ischaemic, BZ = border zone, FAR = remote zone from the infarct. Data are representative of two independent experiments. Each analysed sample was a pool of samples coming from three individuals (*n*=3 animals per group).

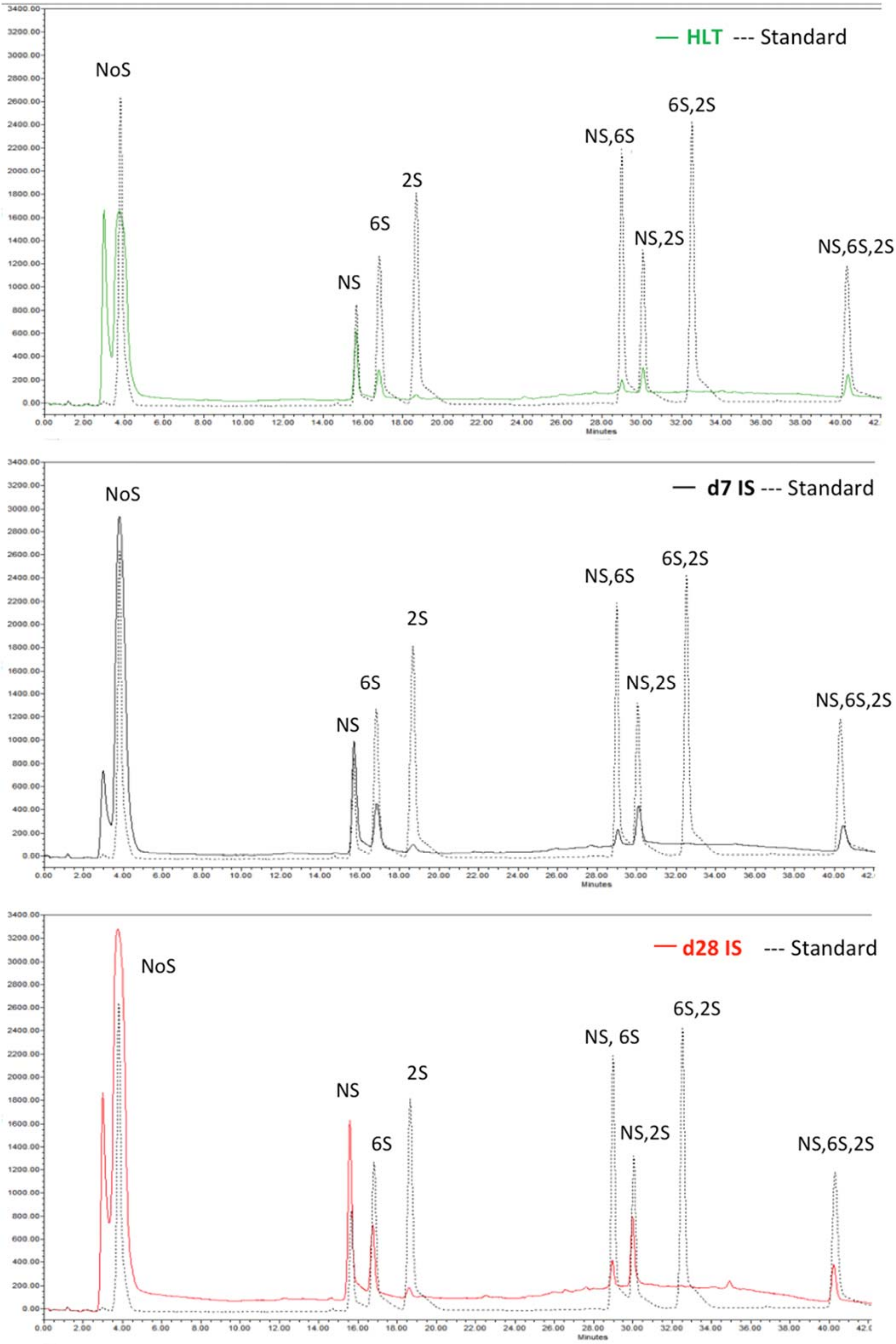

— HLT --- Standard
NoS
6S,2S
NS,6S
2S
6S
NS,2S
NS,6S,2S
NS
Minutes
— d7 IS --- Standard
NoS
NS,6S
6S,2S
2S
6S
NS,2S
NS,6S,2S
NS
Minutes
— d28 IS --- Standard
NoS
6S,2S
NS, 6S
2S
NS
NS,2S
6S
NS,6S,2S
Minutes

**Extended Data Figure 8 | HS sulfation pattern following NSTEMI**

Representative HPLC chromatograms showing the sulfation profile of heparan sulfate (HS) extracted from healthy myocardial tissue (top), and from the ischaemic region at day 7 (middle) and 28 (bottom) post-NSTEMI. Each profile has been integrated by comparison to an external standard mix (line in grey dot in all chromatograms) composed of unsufated disaccharide (NoS), monosulfated disaccharides with a different position of sulfation (NS; 6S; 2S), disulfated disaccharides (NS, 6S; NS,2S; 6S, 2S) and trisulfated disaacharide (NS, 6S, 2S).

**Table 1 | Main upstream regulators from pathway analysis on proteomic data in the ischaemic core 28 days post-NSTEMI**

| Upstream Regulator | Activation z-score | p-value of overlap |
|---|---|---|
| CSF2 | 9,391 | 4,54E-37 |
| IFNg | 7,643 | 2,9E-37 |
| TNF | 7,197 | 1,81E-43 |
| IL1B | 7,063 | 7,25E-26 |
| TGFB1 | 6,737 | 1,35E-50 |
| IL2 | 6,602 | 1,22E-26 |
| NFkB (complex) | 6,509 | 3,19E-18 |
| MYD88 | 6,47 | 9,25E-11 |
| Vegf | 6,417 | 1,32E-16 |
| SMARCA4 | 6,398 | 2,77E-20 |
| EGF | 6,256 | 8,68E-18 |
| SP1 | 6,178 | 1,06E-23 |
| IL1 | 6,121 | 8,6E-15 |
| IL5 | 6,063 | 1,18E-08 |
| FOXM1 | 6,017 | 8,63E-19 |
| AGT | 5,932 | 1,82E-12 |
| GLI1 | 5,772 | 2,49E-12 |
| ERBB2 | 5,672 | 3,25E-22 |
| RAF1 | 5,575 | 6,33E-10 |
| CEBPB | 5,569 | 1,96E-11 |
| STAT4 | 5,47 | 0,0000002 |
| P38 MAPK | 5,468 | 2,05E-16 |
| IGF1 | 5,427 | 9,37E-14 |
| IFNa | 5,351 | 1,19E-11 |
| HGF | 5,312 | 6,74E-12 |
| ARNT2 | 5,292 | 0,0028700 |
| FOXO1 | 5,251 | 8,03E-15 |
| CTNNB1 | 5,157 | 1,41E-18 |
| F2 | 5,134 | 0,0000010 |
| IL6 | 5,13 | 3,22E-32 |

#### Abbreviations

$^{1}$H-NMR: Hydrogen Nuclear Magnetic Resonance

ACN: Acetonitrile

AUC: Area Under The Curve

BSA: Bovine Serum Albumin

BZ: Border Zone

CD: Cluster of Differentiation

CID: Collision-Induced Dissociation

CS: Chondroitin Sulfate

DEG: Differential Expressed Gene

DMMB: Dimethylmethylene Blue Assay

DTT: Dithiothreitol

ECG: Electrocardiogram

ECM: Extracellular Matrix

EDTA: Ethylenediaminetetraacetic Acid
EF: Ejection Fraction
EM: Electron Microscopy
FA: Formic Acid
FAR: Remote Region
FBS: Foetal Bovine Serum
FDR: False Discovery Rate
FGF: Fibroblast Growth Factor
FS: Fractional Shortening
GAGs: Glycosaminoglycans
H&E: Haematoxylin and Eosin
HPLC: High Performance Liquid Chromatography
HS: Heparan Sulfate
IAA: Iodoacetamide
IL: Interleukin
IPA: Ingenuity Pathway Analysis
IS: Ischemic core
LAD: Left Anterior Descending Coronary Artery
MI: Myocardial Infarction
MMP: Matrix Metalloproteinase
MWCO: Molecular Weight Cut-Off
NSTEMI: Non-ST-segment Elevation Myocardial Infarction
O/N: Overnight
PB: Phosphate Buffer
PBS: Phosphate Buffer Saline
PCI: Percutaneous Coronary Intervention
PFA: Paraformaldehyde
PG LC-ESI-MS/MS: Porous Graphite Liquid Chromatography-electrospray Ionization-tandem Mass Spectrometry
PK: Proteinase K
RT: Room Temperature
sGAG: Sulfated Glycosaminoglycan
SMA: Smooth Muscle Actin
STEMI: ST-segment Elevation Myocardial Infarction
TEAB: Triethylammonium Bicarbonate
TEM: Transmission Electron Microscopy
TFA: Trifluoroacetic Acid

TNF: Tumour Necrosis Factor

TTE: Transthoracic Echocardiography

UHPLC: Ultra-High Performance Liquid Chromatography

VEGF: Vascular Endothelial Growth Factor

α-Gal: α-Galactose
